# Pyramidal tract neurons drive feed-forward excitation of striatum through cholinergic interneurons

**DOI:** 10.1101/2020.12.14.422716

**Authors:** Nicolás A. Morgenstern, Ana Filipa Isidro, Inbal Israely, Rui M. Costa

## Abstract

Corticostriatal connectivity is central for many cognitive and motor processes, such as reinforcement or action initiation and invigoration. The cortical input to the striatum arises from two main cortical populations: intratelencephalic (IT) and pyramidal tract (PT) neurons. We uncovered a new feedforward excitatory circuit, supported by a polysynaptic motif from PT neurons to cholinergic interneurons (ChIs) to excitatory inputs, which runs in parallel to the canonical monosynaptic corticostriatal connection. This new motif conveys a delayed second phase of excitation to striatal spiny projection neurons (SPNs), through an acetylcholine-dependent glutamate release mechanism, resulting in biphasic corticostriatal signals. These biphasic signals are a hallmark of PT, but not IT, corticostriatal inputs, due to a stronger relative input from PT neurons to ChIs. These results uncover a novel feed-forward circuit mechanism by which PT activity differentially gates excitatory inputs to the striatum, with potential implications for behavior, plasticity and learning.

**Highlights:**

- PT, but not IT, corticostriatal inputs convey biphasic excitation to SPNs via a disynaptic circuit involving striatal ChIs.
- PT neurons recruit ChIs more efficiently than IT neurons due to a stronger relative input to ChIs.
- The second phase of SPN excitation is mediated by nicotinic receptors at long-range glutamatergic axons in the striatum.
- Suprathreshold depolarizations of SPNs by PT inputs depend on ChI→SPN excitation and result in delayed spiking.

## Introduction

In the brain, the connection from the cortex to the striatum is central for many cognitive and motor processes, such as learning new motor skills or selecting proper actions in response to internal or contextual changes (**Graybiel et al.**, 1994; **Pennartz et al.**, 2009; **Klaus et al.**, 2019). The striatum is the largest input nucleus to the basal ganglia, a group of interconnected subcortical nuclei that regulate brainstem, midbrain and thalamocortical circuits, forming a long-range connectivity loop with the later (**Alexander and Crutcher**, 1990). In the striatum, about 95% of the neurons are GABA-releasing spiny projection neurons (SPNs) (**Kemp and Powell**, 1971; **Gerfen and Wilson**, 1996). Besides being the most abundant ones, SPNs are the only neuronal subtype projecting outside this structure (**Kemp and Powell**, 1971), filtering the information that is outputted to downstream basal ganglia nuclei and, ultimately, modulating brainstem activity, and thalamic and cortical feed-back.

Synaptic input to striatum arises from most cortical areas, and to a lesser extent from the thalamus, through highly organized excitatory long-range axons (**Yeterian and Van Hoesen**, 1978; **Flaherty and Graybiel**, 1993; **Wall et al.**, 2013; **Guo et al.**, 2015; **Hintiryan et al**., 2016; **Hooks et al.**, 2018; **Johansson and Silberberg**, 2020). Given that the striatum lacks intrinsic glutamatergic neurons (**Tepper et al.**, 2007), these inputs onto SPNs and other striatal neuronal subtypes, are key for normal striatal function. Corticostriatal inputs are thought to convey motor and contextual signals to SPNs, information that is critical for proper action selection (**Gremel and Costa**, 2013; **Klaus et al.**, 2019). Corticostriatal contacts are also the site for plasticity underlying striatal-dependent learning (**Kreitzer and Malenka**, 2008; **Lee et al.**, 2015). In fact, synaptic weight changes occur in excitatory contacts onto SPNs when mice learn a motor task (**Yin et al.**, 2009; **Shan et al.**, 2014). Moreover, movement disorders in humans and mouse models of diseases like Parkinson’s affect this connection (**Stephens et al.**, 2005; **Day et al**., 2006; **Fieblinger et al**., 2014; **Hawes et al.**, 2015), highlighting its importance.

The vast majority of the excitatory neurons projecting to the striatum are located in cortical layer (L)5 (**Shepherd**, 2013). However, they are a heterogenous population and could be subdivided into two major categories: intratelencephalic (IT) and pyramidal tract (PT) neurons (**Cowan and Wilson**, 1994; **Molnár and Cheung**, 2006; **Shepherd**, 2013). Each subpopulation has characteristic intracortical laminar position and stereotyped axonal projection patterns, suggesting divergent functionality (**Beloozerova et al.**, 2003; **Graybiel**, 2005; **Reiner**, 2010; **Li et al.**, 2015). The somata of IT neurons span cortical L5 and their axons extend within ipsi- and contralateral cortical areas, and also to the ipsi-and contralateral striatum. On the other hand, PT somata are located in deep L5 and their corticofugal projections send collaterals to several ipsilateral subcortical structures, predominating those to the ipsilateral striatum (**Nelson et al.**, 2020). Moreover, while IT axons synapse onto PTs, direct PT contacts onto ITs are rare, suggesting a hierarchical IT→PT anatomo-functional organization (**Morishima**, 2006; **Brown and Hestrin**, 2009; **Anderson et al.**, 2010; **Kiritani et al.**, 2012). These morphological and connectivity features, together with studies showing that IT and PT have distinct roles in action planning and execution (**Li et al.**, 2015), suggest that ITs are mostly involved in intracortical action preparation, while PTs trigger action execution by broadcasting a command signal throughout the multiple motor-related subcortical structures they innervate.

The striatum is therefore the only non-cortical structure where IT and PT pathways converge (**Shepherd**, 2013), synapsing onto both striatonigral and striatopallidal SPNs (**Kress et al.**, 2013). Thus, understanding the differences between the signals that SPNs receive from these two key cortical afferents, and their impact on striatal output, becomes crucial for tackling neuronal circuits supporting motor learning and behavior.

However, the direct cortex (Cx)→SPN connection is not the sole determinant of striatal output. In fact, SPN spiking is tightly controlled by intrastriatal polysynaptic interactions with sparse local GABAergic and cholinergic interneurons (ChIs)(**Assous and Tepper**, 2018). For instance, parvalbumin-expressing fast-spiking interneurons (FSIs), driven by cortical inputs, exert strong feed-forward inhibition onto SPNs, controlling their output (**Koós and Tepper**, 1999; **Mallet et al.**, 2005; **Gittis et al.**, 2010; **Planert et al.**, 2010). In turn, ChIs regulate striatal function by releasing acetylcholine that acts either through neuromodulatory muscarinic receptors onto SPNs (**Goldberg et al.**, 2012) or through presynaptic nicotinic receptors regulating dopamine, GABA and glutamate release (**Wonnacott et al.**, 2000; **Campos et al.**, 2010; **English et al.**, 2012; **Threlfell et al.**, 2012; **Nelson et al.**, 2014; **Howe et al.**, 2016). Recent studies showed non-uniform and highly specific organization of diverse cortical and thalamic inputs onto striatal interneurons, expanding our knowledge of their afferent connectivity (**Guo et al.**, 2015; **Assous and Tepper**, 2018; **Johansson and Silberberg**, 2020). However, despite their critical influence on SPNs spiking, it is still unknown whether striatal interneurons have biased inputs from IT versus PT neurons.

In this study, we used transgenic mouse lines, optogenetics and slice electrophysiology to investigate the differences between IT and PT corticostriatal connectivity to the striatum. We found a new feedforward excitatory circuit, supported by a polysynaptic motif from PT neurons to ChIs to excitatory inputs, running in parallel to the canonical monosynaptic Cx→SPNs connection. This new motif conveys a delayed second phase of excitation to SPNs, through an acetylcholine-dependent glutamate release mechanism, resulting in biphasic corticostriatal signals. Moreover, we found that these biphasic signals are a hallmark of PT, but not IT, corticostriatal inputs, due to their stronger relative input to ChIs. This work uncovers a novel circuit mechanism by which PT, but not IT neurons, drive feed-forward excitation to the striatum, with potential implications for behavior, plasticity and learning.

## Results

### PT corticostriatal inputs evoke biphasic responses onto SPNs

To investigate the differences between the signals that IT and PT corticostriatal neurons convey to SPNs, we crossed a transgenic mouse line expressing ChannelRhodopsin-2 (ChR2)-EYFP under the control of cre-recombianse (Ai32; *Rosa-CAG-LSL-ChR2(H134R)-EYFP-WPRE)* with a mouse line expressing cre-recombinase in either IT (Tlx3; *Tg(Tlx3-cre)PL58Gsat/Mmucd)* or PT (OE25; *Tg(Chrna2-cre)OE25Gsat/Mmucd)* cortical neurons. This resulted in the selective expression of ChR2-EYFP in one of these two neuronal populations (IT-ChR2-EYFP and PT-ChR2-EYFP, respectively), confirming their laminar location in cortical L5, as well as their long-range axonal projections into the dorsolateral striatum (DLS, **Figure 1A and B**). We then used whole-cell patch-clamp to record excitatory post-synaptic currents (EPSCs) from SPNs in the DLS of acute brain coronal slices while wide-field photostimulating pathway-specific corticostriatal fibers (**Figure 1C**). This is a well-established approach to study long-range connectivity in vitro, because ChR2-expressing axons remain photoexcitable despite losing branches or the connection to their parental soma during the slicing process (**Petreanu et al.**, 2007, 2009; **Yang et al.**, 2013; **D’Souza et al.**, 2016; **Morgenstern et al.**, 2016; **Tanimura et al.**, 2019). As expected, presynaptic photostimulation of identical power but different durations elicited postsynaptic responses of variable amplitude (**Figure 1D**). Surprisingly, IT and PT activation elicited responses with different characteristics. The stimulation of IT fibers evoked typical monophasic EPSCs, consistent with direct excitatory inputs from IT neurons. In turn, PT axons activation often elicited EPSCs with two distinguishable phases, suggesting an additional disynaptic component (**Figure 1D**). Accordingly, when comparing IT→SPN with PT→SPN EPSCs from individual trials showing first peaks of similar amplitude, the overall charge transferred to postsynaptic SPNs was larger for PT→SPNs (**Figure 1E**). To investigate if such charge difference was due to the higher probability of evoking a second peak in PT→SPN EPSCs, we used a threshold to detect the delayed phase of corticostriatal signals (**Methods**). Indeed, the probability of evoking a second phase on SPN EPSCs was higher for PT than for IT stimulation across all first peak amplitudes explored (**Figure 1F**). Moreover, both the amplitude and the charge of the second phase increased with the increase in amplitude of the first peak (**Figures 1G and H**). We standardized comparisons to the amplitude of the EPSC first peak rather than to the phostostimulation conditions because of the variable expression levels of ChR2 across slices, subjects and transgenic mouse lines. However, although PT fibers required, on average, longer illumination times than IT axons for evoking responses within the explored range (IT, 6.05 ± 0.28 ms, range: 0.1 – 30, n= 423; PT, 9.24 ± 0.17 ms, range: 0.3 – 22, n= 797; p=1.01 x 10^-37^, Wilcoxon rank sum test, z=-12.84), when comparing similar photostimulation conditions, the probability of evoking biphasic EPSCs was higher for PT inputs (**Figure 1I**).

**Figure 1.**
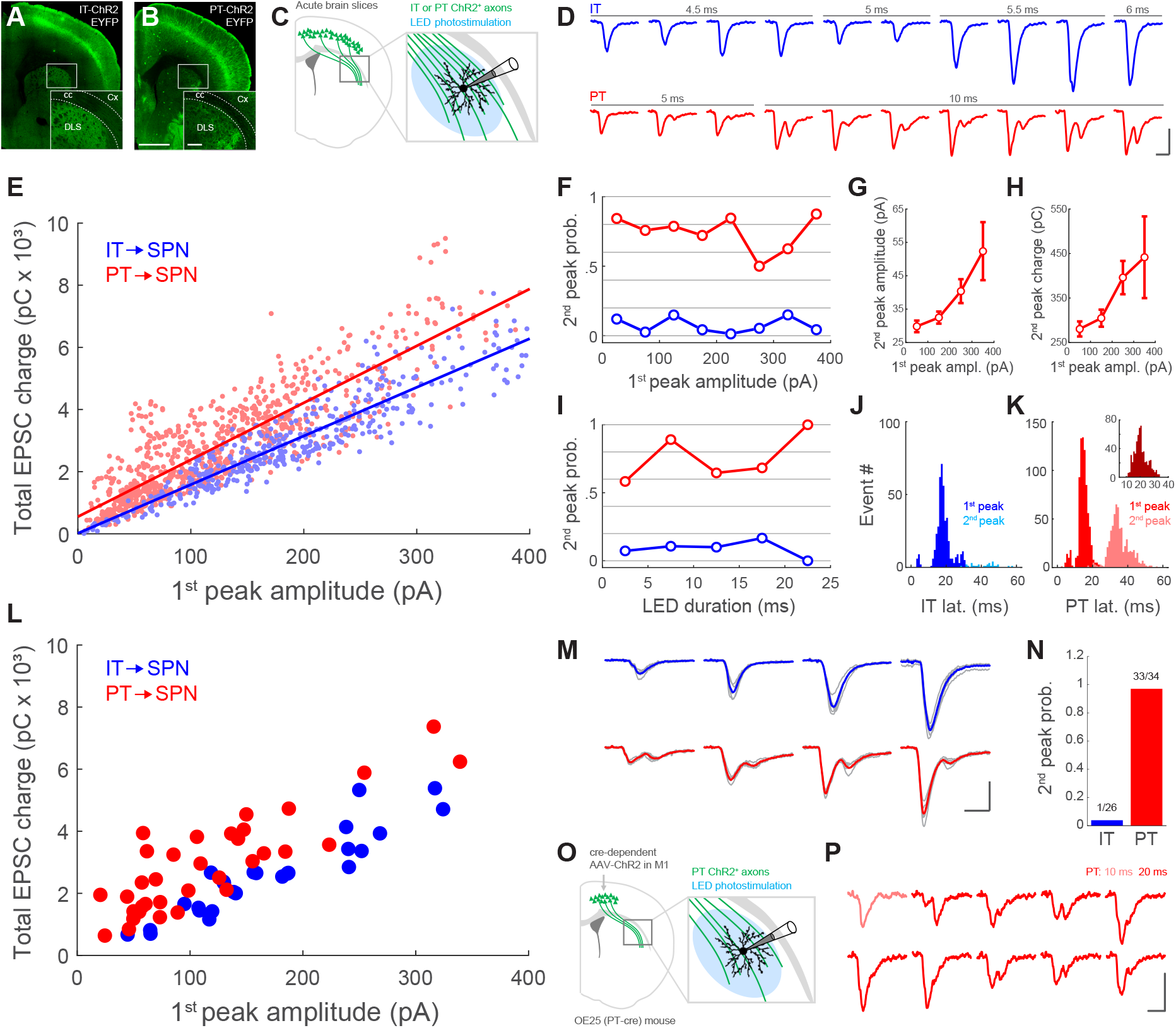
PT corticostriatal inputs evoke biphasic responses onto SPNs. (A, B) 10x confocal image tiles of coronal brain slices from IT-ChR2-EYFP (A) or PT-ChR2-EYFP (B) mice. Scale bar: 1 mm. Insets: magnification of the area in the white boxes. Scale bars: 200 μm. cc: corpus callosum. Cx: cortex. (C) Diagram of experimental design: SPNs from the DLS of IT- or PT-ChR2-EYFP expressing mouse lines were recorded using whole-cell patch-clamp while wide-field photostimulating with LED blue light. (D) Example of voltage-clamp recordings from SPNs when photostimulating IT or PT corticostriatal axons with identical illumination power but different stimulus duration, inter-stimulus interval: 30 s. Scale bar: 20 ms, 100 pA. (E) Total charge of the EPSC as a function of the amplitude of its 1^st^ peak. Each point represents an individual trial. n=423 trials from 26 SPNs from 10 IT-ChR2-EYFP mice (blue). n=797 trials from 34 SPNs from 19 PT-ChR2-EYFP mice (red). Solid lines are a linear fit for each group. Slope with 95% confidence interval (CI): IT, 15.66 pC/pA (15.05, 16.28); PT, 18.32 pC/pA (17.59, 19.06). Intersect with 95% CI: IT, 15.43 pC (−110.1, 141); PT, 550 pC (440.6, 659.5). (F) Probability of evoking a 2^nd^ peak as a function of the 1^st^ peak amplitude when photostimulating IT (blue) or PT (red) axons. Data is binned in 8 bins of 50 pA. (G, H) Mean amplitude (G) and charge (H) of 2^nd^ PT→SPN EPSC peak as a function of the amplitude of the 1^st^ peak. Data is binned in 4 bins of 100 pA. Datapoints are mean ± SEM. (I) Probability of evoking a 2^nd^ peak as a function of photostimulating duration for IT (blue) and PT (red) axons. Data is binned in 5 bins of 5 ms. (J,K) Histogram of the latencies from the illumination start to the 1^st^ or 2^nd^ peak in IT-ChR2-EYFP (J) and PT-ChR2-EYFP (K) mice. IT: n=423 trials from 26 SPNs from 10 mice. PT: n=797 trials from 34 SPNs from 19 mice. Inset in K: histogram showing the latencies from the 1^st^ peak to the 2^nd^ peak in the same trial. n=610 trials from 34 SPNs from 19 PT-ChR2-EYFP mice. (L) Total charge of the mean EPSC for each SPN as a function of the amplitude of its 1^st^ peak. Each point represents an individual neuron. n=26 SPNs from 10 IT-ChR2-EYFP mice (blue). n=34 SPNs from 19 PT-ChR2-EYFP mice (red). (M) Example traces from individual SPNs showing 1^st^ peaks of different amplitudes when photostimulating IT (blue) or PT (red) fibers. Light gray traces are the five individual trials corresponding to the blue/red average traces. Scale bar: 20 ms, 100 pA. (N) Probability of evoking a 2^nd^ peak upon photostimulation of IT (blue) or PT (red) fibers, for individual neurons. n=26 SPNs from 10 IT-ChR2-EYFP mice. n=34 SPNs from 19 PT-ChR2-EYFP mice. (O) Diagram of experimental design: an OE25 (PT-cre) mouse was injected with AAV5-EF1a-DIO-hChR2(H134R)-EYFP-WPRE-pA in the motor cortex (M1). Five weeks later, SPN EPSCs were recorded upon wide-field photostimulation of M1→DLS PT fibers with LED blue light. (P) Example traces from a SPN when photostimulating M1→DLS PT fibers with identical power but different stimulus duration (stimulus duration is color as depicted, inter-stimulus interval: 30 s). Scale bar: 20 ms, 50 pA.

To better characterize the differences between IT and PT corticostriatal signals, we then measured the latencies from the start of illumination to the first and second peak of the EPSCs (**Figures 1J and K**). We found that the overall latency to the first peak was shorter for PT→SPNs when compared to IT→SPNs (PT: 14.91 ± 0.14 ms, n=797 trials; IT: 18.44 ± 0.24 ms, n=423 trials; p=3.99 x 10^-60^, Wilcoxon rank sum test, z=16.36). Moreover, this analysis showed a mean delay of 20.44 ± 0.19 ms (n=610 tirals) from the first to the second peak evoked by PT photostimulation (**Figure 1K, inset**).

We next investigated how the findings from this dataset, displayed above in a trial-by-trial basis, were reflected at the level of individual neurons. Consistently, we found that, when comparing EPSCs in a similar range of first peak amplitudes, SPNs receive more charge from PT than from IT presynaptic neurons (**Figure 1L**). For this analysis, we first averaged five consecutives photostimulation trials, resulting in a mean EPSC for each individual SPN (**Figure 1M**). Furthermore, this approach allowed us to calculate the neuron-based EPSC second peak response probability. In line with the results above, we found a very low probability for evoking biphasic responses when activating IT fibers, but a highly reliable occurrence of EPSC second phases when stimulating PT inputs (**Figure 1N**).

We then designed an experiment to rule out the possibility that the EPSC second phase is elicited by ChR2-expressing long-range axons not originated in the cortex, resulting from extracortical expression of cre-recombinase in the PT-cre OE25 mouse line (**www.gensat.org**). For this purpose, we injected a cre-dependent AAV in the motor cortex of an OE25 mouse, restricting the expression of ChR2 to PT neurons in this area. Five weeks later, we recorded from a SPN in the DLS (**Figure 1O**) and found that exclusive photostimulation of PT cortical axons also evoked biphasic responses in the striatum (**Figure 1P**).

In summary, we found that IT and PT corticostriatal signals are different, with IT inputs evoking mostly monophasic responses, and PT inputs reliably eliciting two sequential excitatory signals that result in biphasic EPSCs onto SPNs.

### Corticostriatal PT→SPN EPSC second peak is mediated by striatal ChIs

In light of our findings, we hypothesized that whereas the first IT and PT EPSC peak is mediated by direct cortical input to SPNs, the second phase reflects intrastriatal polysynaptic feed-forward excitation, preferentially recruited by PT long-range axons. We dismissed a potential contribution of intracortical polysynaptic interactions, because most cortical axons innervating the DLS are severed from their somas upon coronal slice preparation, due to the anatomy of cortical projections. In addition, focal presynaptic phostostimulation with a laser beam in the near vicinity of the recorded SPNs reliably evoked biphasic PT→SPN EPSCs (**Supplemental Figure 1A**), further supporting a role for local intrastriatal interactions in the late phase of the EPSC.

More than 95% of striatal neurons are GABAergic SPNs (**Kemp and Powell**, 1971; **Gerfen and Wilson**, 1996). Moreover, with exception of a small population of acetylcholine-releasing ChIs, all other local interneurons also release GABA (**Assous and Tepper**, 2018). Since in our experimental conditions SPNs were clamped at a membrane potential below the chloride reversal potential (SPN Vholding=-80 mV, Ecl = −75.6 mV), GABA receptor activation would result in excitatory rather than inhibitory currents (**Plenz**, 2003). For this reason, we first investigated whether GABAergic neurotransmission is underlying the second phase of PT→SPN excitation. For this purpose, we photostimulated PT axons while monitoring SPN EPSCs in the absence or presence of the GABA_A_ receptor antagonist picrotoxin (PTX). Since the charge and amplitude of the EPSC second phase increase with the amplitude of the first peak (**Figure 1G and H**), we calculated the charge ratio and the peak ratio to relate the magnitude of both phases (**Methods**). These normalized metrics, as well as the second peak response probability, resulted robust to fluctuations in the amplitude of the first peak (**Supplemental Figure 2**), allowing us to make comparisons across trials within the same SPN, as well as across different SPNs. Strikingly, the addition of PTX to the extracellular solution had no effect on the charge ratio (**Figures 2A, B and C**). Even though PTX elicited a moderate decrease in the peak ratio (**Figure 2D**) the magnitude of such decrease was not enough to fully suppress the second phase of EPSCs in any of the recorded SPNs (**Figures 2B and K**). These data suggest little involvement of GABAergic transmission in the second phase of PT→SPN responses. In addition, the PT→SPN second phase did not change its polarity when the postsynaptic membrane was clamped above the chloride reversal potential (**Supplemental Figure 1B**), further supporting that it is not mediated by GABA.

**Figure 2.**
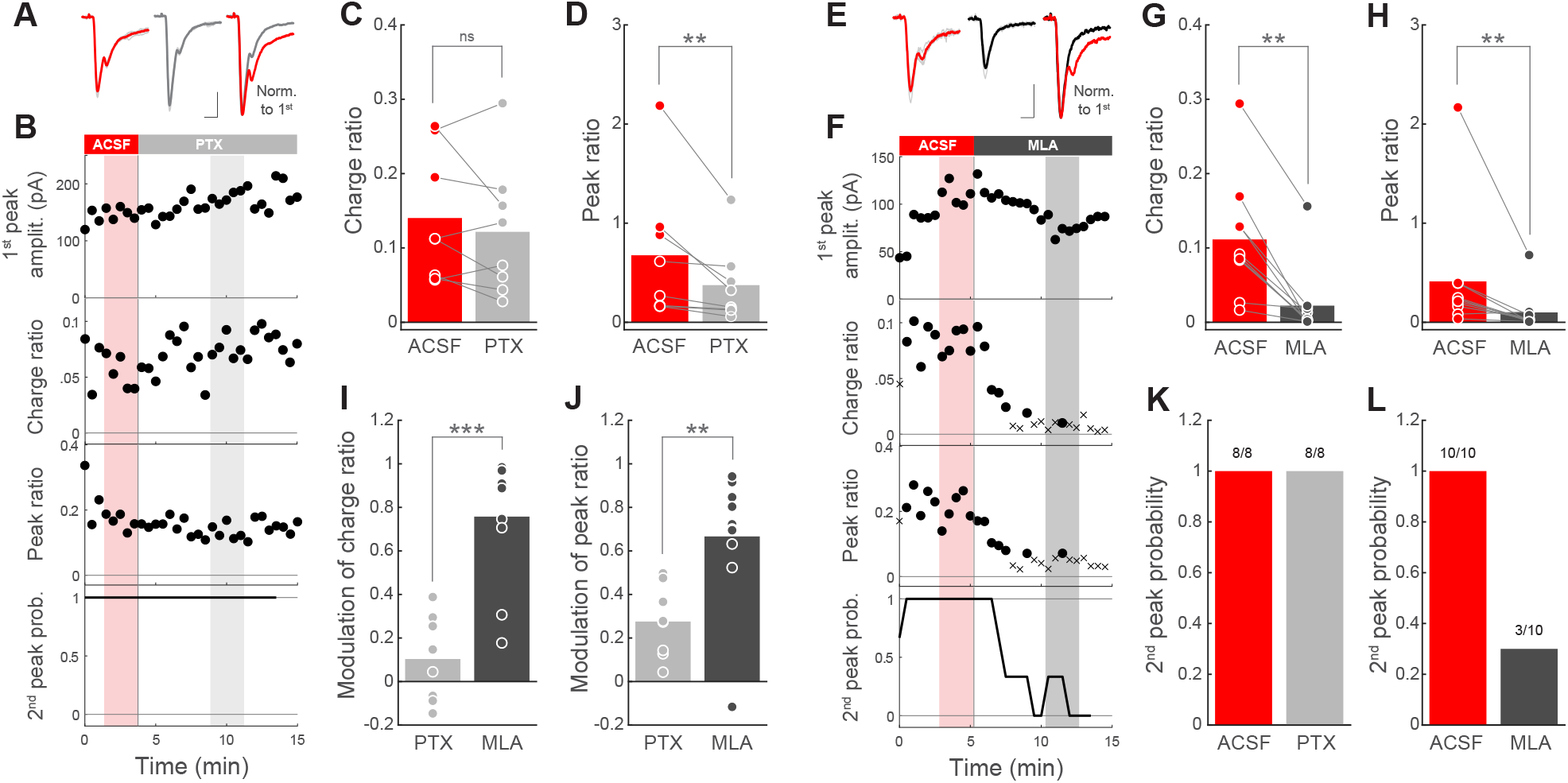
Corticostriatal PT→SPN EPSC second peak is mediated by striatal ChIs. (A) Left, EPSCs from a representative SPN when identically photostimulating PT fibers in ACSF (red) or PTX (gray). Thin light gray traces are the five individual trials corresponding to the thicker mean traces. Scale bar: 20 ms, 50 pA. Right, mean traces normalized to the first peak. (B) From top to bottom: plots showing the amplitude of the 1^st^ peak of the EPSCs, the charge ratio, the peak ratio and the response probability for a 2^nd^ peak as a function of time, for the whole experiment in A. Data points are individual trials elicited every 30 s (for charge ratio and peak ratio, filled circles are EPSCs with a detected 2^nd^ peak). Response probability was calculated within a moving window of 3 consecutive trials. Shadowed areas highlight the individual trials (thin light gray traces in A) averaged for each condition in A (red: ACSF; gray: PTX). (C, D) Charge ratio (C) and peak ratio (D) for individual SPNs in ACSF (red) and PTX (gray) conditions. n=8 SPNs from 5 PT-ChR2-EYFP mice. Bars represent mean. p=0.54688 (C); p=0.0078125 (D); Wilcoxon signed-rank test. (E) Left, EPSCs from a representative SPN when identically photostimulating PT fibers in ACSF (red) or MLA (black). Thin light gray traces are the five individual trials corresponding to the thicker mean traces. Scale bar: 20 ms, 50 pA. Right, mean traces normalized to the first peak. (F) From top to bottom: plots showing the amplitude of the 1^st^ peak of the EPSCs, the charge ratio, the peak ratio and the response probability for a 2^nd^ peak as a function of time, for the whole experiment in E. Data points are individual trials elicited every 30 s (for charge ratio and peak ratio, circles: EPSC with 2^nd^ peak; crosses: EPSC without 2^nd^ peak). Response probability was calculated within a moving window of 3 consecutive trials. Shadowed areas highlight the individual trials (thin light gray traces in E) averaged for each condition in E (red: ACSF; black: MLA). (G, H) Charge ratio (G) and peak ratio (H) for individual SPNs in ACSF (red) and MLA (black) conditions. n=10 SPNs from 7 PT-ChR2-EYFP mice. Bars represent mean. p=0.0019531 (G); p=0.0039063 (H); Wilcoxon signed-rank test. (I, J) Modulation index (**Methods**) of charge ratio (I) and peak ratio (J) for individual SPNs in PTX (gray) or MLA (black) conditions. PTX, n=8 SPNs from 5 PT-ChR2-EYFP mice; MLA, n=10 SPNs from 7 PT-ChR2-EYFP mice. Bars represent mean. p=0.00054847 (I); p=0.0030623 (J); Wilcoxon rank sum test. (K, L) 2^nd^ peak response probability upon photostimulation of PT fibers for individual SPNs before (red) and after the addition of PTX (gray, K) or MLA (black, L). PTX, n=8 SPNs from 5 PT-ChR2-EYFP mice; MLA, n=10 SPNs from 7 PT-ChR2-EYFP mice.

Having ruled out the role of GABAergic interneurons, we postulated that ChIs could mediate these local polysynaptic responses triggered by PT. In fact, current-voltage curves of PT→SPN second phase resembled cationic currents compatible with glutamate or acetylcholine acting through nicotinic receptors (**Supplemental Figure 1C and D**). To test this, we recorded SPNs while photostimulating PT axons without or with the nicotinic acetylcholine receptor antagonist methyllycaconitine (MLA) in the bath. MLA significantly reduced the charge ratio, the peak ratio and the second peak response probability of PT→SPN responses (**Figures 2E-H**), which partially recovered after MLA wash out (**Supplemental Figures 2B and C**). Notably, MLA preferentially impacted on the second, rather than on the first, phase of these EPSCs (**Supplemental Figures 2 and 3**), indicating that it did not reduce overall excitability. Moreover, when comparing the magnitude of the reduction exerted by MLA or PTX, we found a significantly stronger effect of the cholinergic blocker on both the charge ratio and the peak ratio (**Figures 2I and J**). Furthermore, MLA reduced the PT→SPN EPSC second phase to undetectable levels in most of the recorded neurons (**Figure 2L**), implicating ChIs in these delayed signals. In summary, our data strongly suggests that the second excitatory component of the biphasic PT→SPN responses is mediated by nicotinic receptors that are activated upon the selective recruitment of local striatal ChIs by PT input stimulation.

### PT neurons provide stronger relative inputs to ChIs than IT neurons

In the context of our results, we wondered if different connectivity rules operating for each corticostriatal pathway could explain the preferential recruitment of ChI→SPN excitation by PT, but not by IT activation. In that sense, we hypothesized that the relative input strength to different striatal postsynaptic neuronal subtypes is differentially balanced for the IT and PT pathways. Thus, we designed an experiment to test whether cortical IT and PT neurons contact striatal ChIs monosynaptically, and if so, how the connection strength from each pathway is distributed between ChIs and SPNs.

In brains slices of IT- or PT-ChR2-EYFP mice, we first patched a ChI in the DLS (**Figure 3A and B**). We initially recorded visually identified putative ChIs and confirmed their cholinergic identity by assessing their typical electrophysiological properties (depolarized resting membrane potential, presence of sag upon hyperpolarization, spike half-width >1 ms and regular spiking; **Figure 3C and Supplemental Figure 4**) in drug-free extracellular solution. We further confirmed the cholinergic phenotype of the recorded neurons by their characteristic larger somatic area (ChIs: 239.86 ± 11.79 μm^2^, range: 146.32 – 356.6, n=19; SPNs: 108.46 ± 6.53 μm^2^, range: 59.57 – 160.62, n=20; p=1.6 x 10^-7^, Wilcoxon rank sum test, z=-5.24, **Figure 3B**) and their reactivity to Choline-Acetyl Transferase (ChAT) immunolabelling (16 out of 19 ChIs were recovered and identified as ChAT^+^, **Figure 3C**).

**Figure 3.**
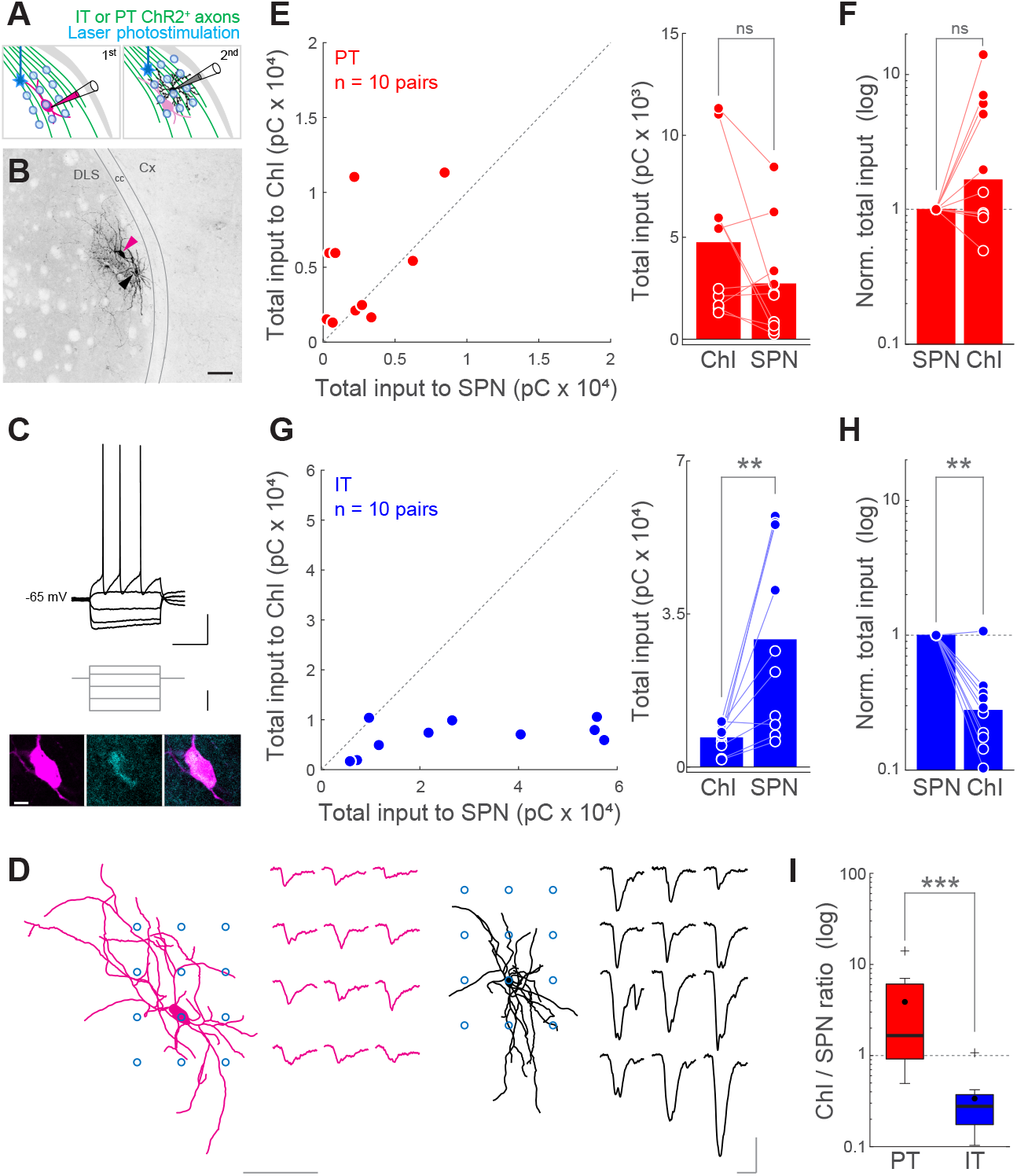
PT neurons provide stronger relative inputs to ChIs than IT neurons. (A) Diagram of experimental design. Left: first, a ChI from the DLS was recorded while IT- or PT-ChR2^+^ presynaptic terminals were photostimulated following a 3 x 4 grid pattern using a 1-photon blue laser in the presence of TTX and 4-AP. Right: second, a neighboring SPN (< 100 μm apart) was recorded while photostimulating with the same grid pattern at identical illumination conditions. (B) 25x confocal image showing a pair of ChI (magenta arrowhead) and SPN (black arrowhead) in the DLS of an IT-ChR2-EYFP mouse filled with biocytin during the recordings and revealed with streptavidin. Scale bar: 100 μm. cc: corpus callosum. Cx: cortex. (C) Top: Input-output curve showing the firing pattern of the ChI in B, scale bar: 20 mV, 500 ms. Gray traces are current injection steps; scale bar: 100 pA. Bottom: 20x single confocal planes showing the neuronal body of the ChI in B (left, magenta), its immunoreactivity to anti-ChAT antibodies (center, cyan) and the overlap of both signals (right). Scale bar: 10 μm. (D) Left, reconstruction of the neuronal morphology from the ChI in B with the relative position of its photostimulation grid (blue circles) and the respective 3 x 4 matrix of recorded EPSCs (magenta traces). Right, reconstruction of the neuronal morphology from the SPN in B with the relative position of its photostimulation grid (blue circles) and its correspondent 3 x 4 matrix of recorded EPSCs (black traces). Left scale bar: 100 μm. Right scale bar: 20 ms, 50 pA. (E) Comparison of total input charge transferred to pairs of neighboring ChIs and SPNs when activating PT corticostriatal inputs. Left, each point represents a pair. Right, each point represents a neuron. Bars represent mean. p=0.16016; n=10 pairs from 9 mice; Wilcoxon signed-rank test. (F) Total input to ChI-SPN pairs normalized to the total input to the SPN, when activating PT fibers. Bars represent median. PT→SPN vs. PT→ChI, p=0.10547; Wilcoxon signed-rank test. n=10 pairs from 9 mice. (G) Comparison of total input charge transferred to pairs of neighboring ChIs and SPNs when activating IT corticostriatal inputs. Left, each point represents a pair. Right, each point represents a neuron. Bars represent mean. p=0.0039063; n=10 pairs from 10 mice; Wilcoxon signed-rank test. (H) Total input to ChI-SPN pairs normalized to the total input to the SPN, when activating IT fibers. Bars represent median. IT→SPN vs. IT→ChI, p=0.0039063; Wilcoxon signed-rank test. n=10 pairs from 10 mice. (I) Box plot comparing the normalized input to ChIs (ChI/SPN ratio) from PT (red) and IT (blue). p=0.00058284; Wilcoxon rank sum test, z=3.44. PT: n=10 pairs from 9 mice; IT: n=10 pairs from 10 mice. Horizontal black line: median; filled black circle: mean; box edges: 25^th^ and 75^th^ percentile; whiskers: maximum and minimum excluding outliers; crosses: outliers.

After that, we added 4-aminopiridine (4-AP) and tetradotoxine (TTX) to the extracellular solution. This drug combination suppresses spiking while preserving ChR2^+^ synaptic terminals ability to release neurotransmitter when photostimulated, thus, restricting EPSCs only to monosynaptic contacts (**Petreanu et al.**, 2009; **Yang et al.**, 2013; **D’Souza et al.**, 2016; **Morgenstern et al.**, 2016). In these conditions, we photostimulated IT or PT corticostriatal presynapses with a blue laser following a grid pattern over the dendrites of the postsynaptic ChI (**Figure 3D**). For each neuron, we computed the total input charge by summing all EPSCs. Since both IT and PT corticostriatal pathways stimulation elicited monosynaptic responses on ChIs, we calibrated the illumination conditions to evoke comparable IT→ChI and PT→ChI EPSCs across experiments (IT: 6.77 ± 1.01 pC*10^3^, range: 1.69 – 10.57, n=10; PT: 4.76 ± 1.34 pC*10^3^, range: 1.31 – 11.32, n=9; p=0.2775, Wilcoxon rank sum test). Ideally, we would have recorded first from SPNs and calibrated photostimulation to evoke similar IT→SPN and PT→SPN EPSCs. However, the need for confirming ChIs electrophysiogical phenotype in the absence of drugs, forced us to record first from ChIs. Right after assessing inputs to the ChI, we recorded a neighboring SPN (less than 100 μm apart, **Figure 3A, B and Supplemental Figure 4E**) and repeated the photostimulation protocol with identical illumination conditions (**Figure 3D**). Since neighboring neurons could potentially sample from the same population of afferent axons, with this experimental design we could compare the pathway-specific input strength balance to ChI-SPN pairs, irrespective of ChR2 expression level variations across slices (**Petreanu et al.**, 2009; **Yang et al.**, 2013; **D’Souza et al.**, 2016; **Morgenstern et al.**, 2016).

We found that the input strength is differentially weighted for each corticostriatal pathway. While PT inputs exhibited similar strengths across ChIs and SPNs (**Figure 3E and F**), the IT connection was significantly biased towards the SPNs (**Figure 3G and H**). To further quantify how the connection strength of each pathway distributes between ChIs and SPNs, we normalized the corticostriatal input that a given ChI-SPN pair receive to the input of that SPN (**Figure 3F and H**). We found that the PT→ChI relative input strength is variable, with a population median of 1.66 times the PT→SPN input (**Figure 3F**). Contrarily, IT→ChI relative input strength was consistently weaker, with a population median of 0.28 times the IT→SPN input (**Figure 3H**). These data strongly suggest that IT and PT pathways follow different connectivity rules when synapsing onto striatal neurons. Moreover, the normalized PT→ChI input was one order of magnitude stronger than the normalized IT→ChI input (**Figure 3I**), supporting a model where, provided similar input to SPNs, PT neurons are more likely to recruit ChIs than IT neurons. Importantly, these differences in IT and PT connectivity could not be explained by differences in the size of the dendrites of the recorded ChIs (**Supplemental Figure 4G**) nor by the distance between ChIs and SPNs within the same pair (**Supplemental Figure 4E**).

Our findings support the co-existence of two parallel excitatory corticostriatal circuits. On one hand, there is the canonical direct long-range connection from cortical neurons to SPNs, which photoactivation elicits the monosynaptic first phase of the IT→SPN and PT→SPN EPSCs. On the other hand, we described a novel parallel direct long-range projection to striatal ChIs from both, IT and PT neurons, but with higher relative input strength for PT→ChIs than for IT→ChIs. As a consequence, only PT inputs effectively recruit ChIs upon photostimulations that evoke similar EPSC first peak amplitudes onto SPNs. Together with our previous data, these results support a corticostriatal Cx→ChIs→SPNs feed-forward excitatory motif, and explain why PT inputs are more likely than IT inputs to evoke biphasic corticostriatal EPSCs onto SPNs.

### ChIs indirectly excite SPNs via an acetylcholine-dependent glutamate release mechanism

We next wanted to dissect the precise circuit mechanism underlying ChI→SPN EPSCs. One possibility is that ChIs form direct monosynaptic contacts onto SPNs supporting nicotinic transmission, which existence, to our understanding, is not described in the literature (**Zhou et al.**, 2002; **Assous and Tepper**, 2018). Alternatively, ChIs could indirectly convey signals to SPNs by nicotinic activation of presynaptic glutamatergic axons (**Campos et al.**, 2010; **Howe et al.**, 2016; **Abudukeyoumu et al.**, 2018). Interestingly, this interaction between cholinergic and glutamatergic systems is mediated by α7-nicotinc receptors, which are a major target of MLA, the cholinergic antagonist proved to suppress the second phase of PT→SPN EPSCs (**Figure 2**).

To investigate these two possibilities, we used a double transgenic mouse line *(ChAT-Cre x Ai32)* where ChR2 is expressed in acetylcholine-releasing neurons (ChAT-ChR2-EYFP). Since the cholinergic innervation of the striatum is dominated by local ChIs, rather than extrinsic innervation (**English et al.**, 2012; **Goldberg et al.**, 2012; **Dautan et al.**, 2014; **Deffains and Bergman**, 2015), we reasoned that photostimulation in this preparation would mostly reflect the activation of striatal ChIs (**Figure 4A**).

**Figure 4.**
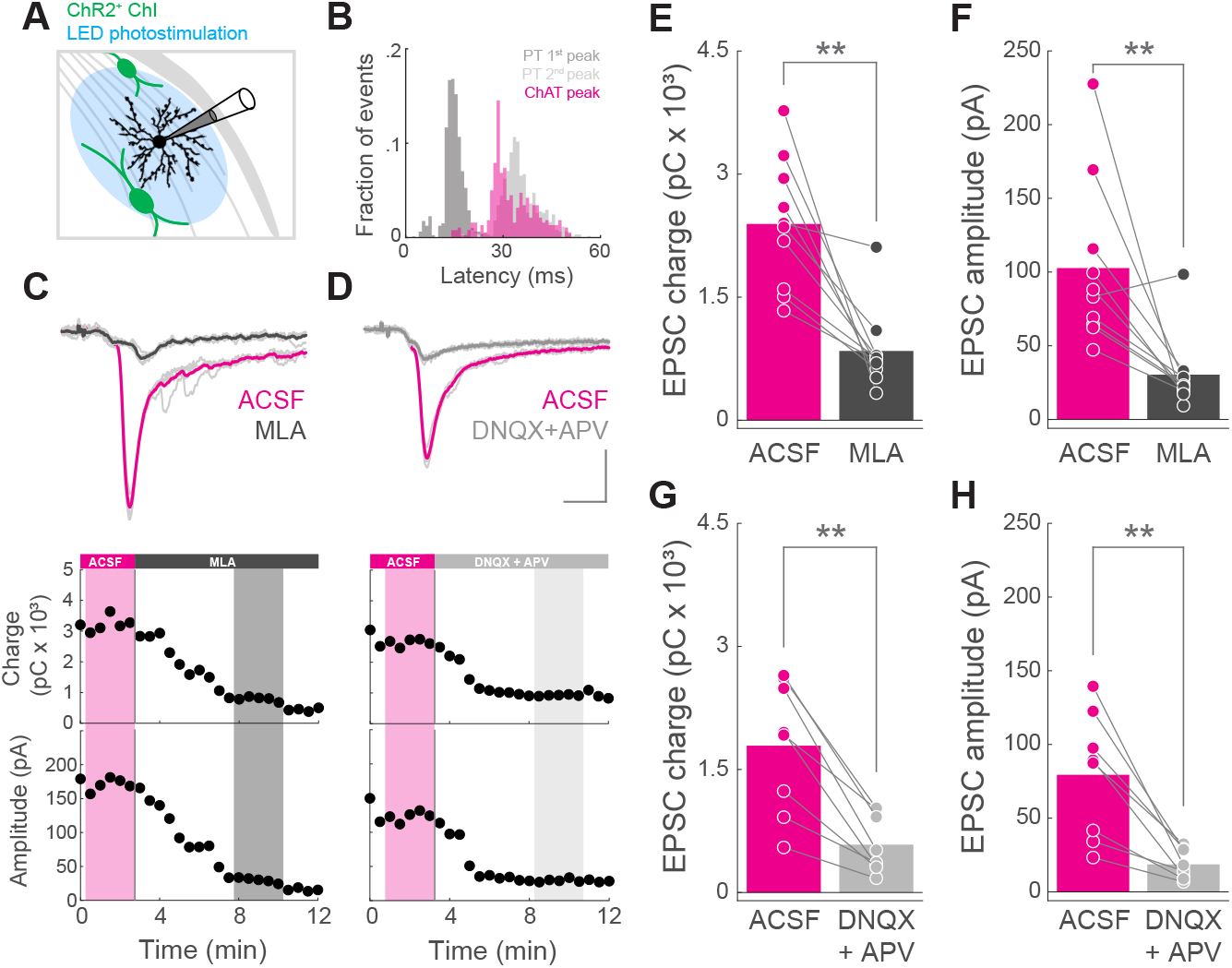
ChIs indirectly excite SPNs via an acetylcholine-dependent glutamate release mechanism. (A) Diagram of experimental design: SPNs from the DLS of ChAT-ChR2-EYFP mice were recorded using whole-cell patch-clamp while field photostimulating with LED blue light. (B) Histogram showing the latencies from the start of illumination to the peak of the EPSC in the SPNs when photostimulating ChIs (magenta). n=379 trials from 20 SPNs from 9 ChAT-ChR2-EYFP mice. For comparison, the distribution of the latencies to the 1^st^ (dark gray) or 2^nd^ peak (light gray) in PT-ChR2-EYFP mice (same data than Figure 1K) is shown. (C) Top: example of an individual experiment showing EPSCs from a SPN when identically photostimulating ChAT-ChR2-EYFP neurons before (ACSF, magenta) and after the application of MLA (black). Thin light gray traces are individual trials corresponding to the thicker mean traces. Same scale as in D. Bottom: plots showing the EPSC charge and peak amplitude as a function of time, for the same experiment of the traces above. Black filled circles represent individual trials elicited every 30 s. Shadowed areas highlight the individual trials (thin light gray traces above) averaged for each condition (magenta: ACSF; dark gray: MLA). (D) Top: example of an individual experiment showing EPSCs from a SPN when identically photostimulating ChAT-ChR2-EYFP neurons before (ACSF, magenta) and after the application of DNQX and APV (gray). Thin light gray traces are individual trials corresponding to the thicker mean traces. Scale bar: 25 ms, 50 pA. Bottom: plots showing the EPSC charge and peak amplitude as a function of time, for the same experiment of the traces above. Black filled circles represent individual trials elicited every 30 s. Shadowed areas highlight the individual trials averaged for each condition (magenta: ACSF; light gray: DNQX and APV). (E, F) EPSC charge (E) and amplitude (F) for individual SPNs in ACSF (magenta) and MLA (black) conditions. n=10 SPNs from 5 ChAT-ChR2-EYFP mice. Bars represent mean. p=0.0019531 (E); p=0.0039063 (F); Wilcoxon signed-rank test. (G, H) EPSC charge (G) and amplitude (H) for individual SPNs in ACSF (magenta) and DNQX and APV (gray) conditions. n=8 SPNs from 4 ChAT-ChR2-EYFP mice. Bars represent mean. p=0.0078125 (G); p=0.0078125 (H); Wilcoxon signed-rank test.

We could reliably evoke EPSCs on the recorded SPNs when photoactivating ChIs (**Figure 4C and D**). Strikingly, when compared to the PT→SPN latency to the first peak, the ChI→SPN EPSC peak latency was longer (PT first peak: 14.91 + 0.14 ms, n=797; ChI→SPN peak: 32.33 + 0.37 ms, n=379; Kruskal-Wallis test and multicompare post hoc Dunn’s test, p<0.0001). Indeed, this latency resembled that from PT→SPN second phase (ChI→SPN peak: 32.33 + 0.37 ms, n=379; PT second peak: 35.4 + 0.24 ms, n=610, Kruskal-Wallis test and multicompare post hoc Dunn’s test test, p<0.0001; **Figure 4B**), suggesting a delayed mechanism, more compatible with indirect transmission.

To confirm that these EPSCs were mediated by a similar circuit to PT→SPN EPSC second phase, and not by direct glutamate release from ChIs (**Higley et al.**, 2011), we tested the susceptibility of these responses to MLA. Both the charge and the amplitude of ChI→SPN EPSCs were significantly reduced by MLA (**Figure 4C, E and F**), indicating the necessity of nicotinic activation to excite SPNs.

We then reasoned that if ChI→SPN EPSCs are mediated by monosynaptic contacts; they should be insensitive to the blockade of glutamatergic transmission. If, alternatively, ChIs signals to SPNs involve the recruitment of presynaptic glutamatergic terminals, EPSCs should then be modulated by blocking glutamate receptors. Indeed, the addition of the AMPA and NMDA receptor blockers 6,7-dinitroquinoxaline-2,3-dione (DNQX) and (2R)-amino-5-phosphonovaleric acid (APV), respectively, significantly reduced the charge and amplitude of ChI→SPN EPSCs (**>Figure 4D, G and H**). Altogether, these results support the hypothesis that ChIs mediate the late phase of PT→SPN EPSCs through acetylcholine-dependent glutamate release from excitatory long-range axons innervating DLS.

### Activation of PT corticostriatal inputs evoke delayed SPN spiking

The data presented so far supports a new excitatory long-range corticostriatal circuit consisting of PT→ChIs→glutamatergic axons (glutAx)→SPNs. We next investigated how this feed-forward excitation impacts on SPN spiking, and in turn, on striatal output. We started by photostimulating IT or PT fibers while monitoring the membrane potential of SPNs using current-clamp recordings. Not surprisingly, subthreshold responses from these experiments paralleled our previous findings using voltage-clamp (**Figure 1**). We found that excitatory postsynaptic potentials (EPSPs) of similar first peak amplitude had larger area when elicited by PT activation, both at the trial-by-trial (**Figure 5A**) and at the individual SPN level (**Figure 5B**). Consistently, we found that such area difference is due to biphasic EPSPs (**Figure 5C**), which occur with higher probability when PT, rather than IT, axons are activated (**Figure 5D**).

**Figure 5.**
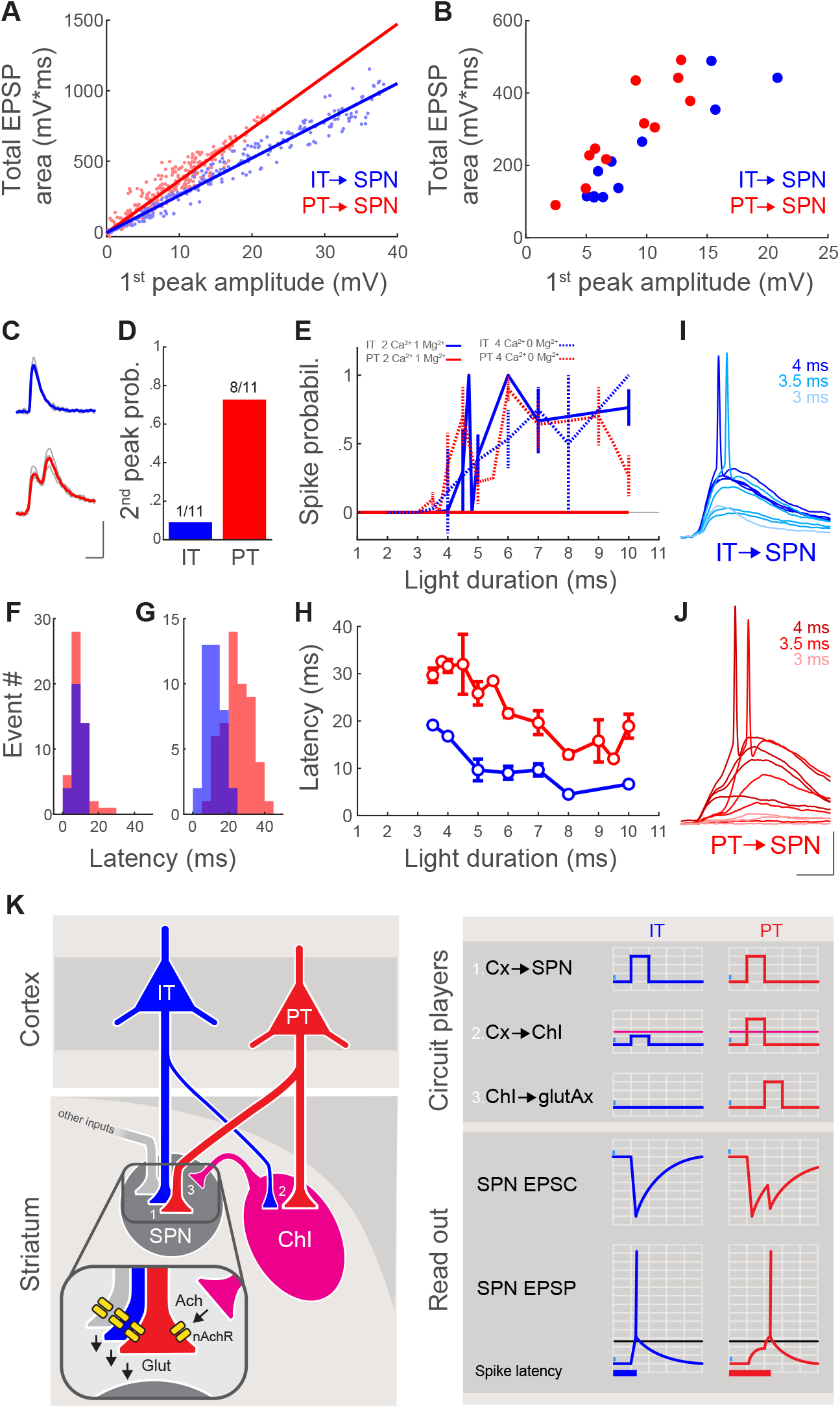
Activation of PT corticostriatal inputs evoke delayed SPNs spiking. (A) Total area of the EPSP evoked upon photostimulation of corticostriatal IT (blue) or PT (red) fibers with LED blue light as a function of the amplitude of its 1^st^ peak. Each data point represents an individual trial. n=286 trials from 11 SPNs from 4 IT-ChR2-EYFP mice (blue); n=384 trials from 11 SPNs from 4 PT-ChR2-EYFP mice (red). Solid lines are a linear fit for each group. Slope with 95% CI: IT, 26.47 pC/mV*ms (25.95, 26.99); PT, 37.05 pC/mV*ms (36.07, 38.03). Intersect with 95% CI: IT, −7.47 pC (17.11, 2.17); PT, −8.68 pC (−17.83, 0.47). (B) Total area of the mean EPSP for each SPN as a function of the amplitude of its 1^st^ peak. Each point represents an individual neuron. n=11 SPNs from 4 IT-ChR2-EYFP mice (blue). n=11 SPNs from 4 PT-ChR2-EYFP mice (red). (C) Example traces from individual SPNs when photostimulating IT (blue) or PT (red) fibers. Light gray traces are the individual trials underlying the blue/red mean traces. Scale bar: 25 ms, 5 mV. (D) Probability of evoking a 2^nd^ EPSP peak upon photostimulation of IT (blue) or PT (red) fibers, for individual neurons. n=11 SPNs from 4 IT-ChR2-EYFP mice. n=11 SPNs from 4 PT-ChR2-EYFP mice. (E) Spike probability as a function of photostimulation duration recorded from SPNs when activating IT (blue) or PT (red) fibers in normal ACSF (solid line) or ACSF with 0 mM Mg^2+^ and 4 mM Ca^2+^ (dotted line). Datapoints are mean ± SEM. (F, G) Histogram showing the latencies from the beginning of the illumination to the start of the EPSP (F) and from the start of the EPSP to the peak of the spike (G) in IT- (blue) or PT-ChR2-EYFP (red) mice recorded in ACSF with 0 mM Mg^2+^ and 4 mM Ca^2+^. p= 0.2975 (F); p=2.28 x 10^-11^ (G); Wilcoxon rank sum test. IT: n=38 spikes from 13 SPNs from 11 mice. PT: n=52 spikes from 7 SPNs from 7 mice. (H) Spike latency as a function of photostimulation duration recorded from SPNs when activating IT (blue) or PT (red) fibers in ACSF with 0 mM Mg^2+^ and 4 mM Ca^2^. Datapoints are mean ± SEM. (I) Example traces recorded from the same SPN when photostimulating IT fibers with increasing stimulus duration in ACSF with 0 mM Mg^2+^ and 4 mM Ca^2+^. Same scale as in J. (J) Example traces recorded from the same SPNs when photostimulating PT fibers with increasing stimulus duration in ACSF with 0 mM Mg^2+^ and 4 mM Ca^2+^. Scale bar: 25 ms, 20 mV. (K) Summary of the results: left, circuit diagram proposed for corticostriatal connectivity. IT and PT cortical neurons project to both SPNs (1) and ChIs (2). While PT→SPNs (1, red) and PT→ChIs (2, red) input strength is similar, IT→ChIs (2, blue) connection is weaker than IT→SPNs (1, blue). Within the striatum, ChIs convey excitation to SPNs by recruiting long-range glutamatergic terminals reaching DLS (3 and inset). Ach, acetylcholine; nAchR, nicotinic receptors; Glut, glutamate. Right, schematic of the activation of the different circuit players upon IT or PT photostimulation and its impact on the recorded SPNs. Magenta and black horizontal lines represent ChIs and SPNs spiking threshold, respectively.

Although in the intact brain SPN spiking would probably depend on simultaneous IT and PT EPSCs, our approach gave us the unique opportunity to independently explore how IT or PT inputs control SPNs spiking, which is key to understanding corticostriatal connectivity in further detail. However, in normal ACSF conditions, only IT activation successfully evoked spiking at SPNs (**Figure 5E**). To overcame this, we reproduced the latter experiments in low-magnesium and high-calcium ACSF (0 mM Mg^2+^, 4 mM Ca^2+^) pursuing the facilitation of polysynaptic activity. In this preparation, IT and PT photostimulation evoked spiking in SPNs with similar probability (**Figure 5E**) but different latency. Overall latency from the start of the illumination until the spike peak revealed significantly longer for PT activation (IT, 20.57 ± 0.58 ms, range: 15.9 – 30.8, n=38; PT, 34.48 ± 1.16 ms, range: 18.2 – 55.1, n=52; p=3.59 x 10^-12^, Wilcoxon rank sum test, z=-6.95). Interestingly, while IT→SPN spike latency resembled the timing to the first EPSC peak, the latency to PT→SPN spikes was similar to the second EPSC peak (**Figure 1J and K**). However, spike latency differences could also emerge from differences in ChR2 expression levels in IT and PT lines. For instance, lower ChR2 expression levels in PT than in IT axons would delay the start of the PT→SPN EPSPs, shifting the peak of the action potentials towards the right. To rule out this possibility, we separately analyzed the latency from the beginning of the illumination until the EPSP start and from this point until the action potential peak (**Methods**). We found that while the latency until the beginning of the EPSP was indistinguishable between IT and PT (**Figure 5F**), PT-triggered spikes developed significantly later after that point (**Figure 5G**). These results indicate that, at least in these conditions, IT and PT presynaptic activation speed is comparable, and rule out the possibility that overall spiking latency differences are due to variable levels of ChR2 expression. Importantly, the overall light durations used for evoking IT- and PT-dependent spiking were similar (IT, 5.95 ± 0.31 ms, range: 3.5 – 10, n=38; PT, 6.13 ± 0.30 ms, range: 3.5 – 10, n=52; p=0.8523, Wilcoxon rank sum test, z= −0.19), and when comparing identical light conditions, the latency from EPSP start until action potential peak was always longer for PT→SPN spikes than for IT→SPN spikes (**Figure 5H**).

Altogether, this data indicates that while IT-triggered spiking occurs by the faster monosynaptic depolarization of SPNs, PT→SPNs direct depolarization is less likely to evoke suprathreshold responses, and spiking threshold is most likely reached during the delayed feed-forward ChIs→glutAx→SPNs excitation phase. This becomes especially evident in conditions that favor polysynaptic activity, and it is reflected in the fact that PT→SPN spikes, in contrast to IT→SPN spikes, occur at the start of the EPSP second phase (**Figure 5I and J**).

## Discussion

Our data uncovers a new circuit mechanism by which IT and PT corticostriatal inputs differentially impact SPNs (**Figure 5K**). We found a feed-forward corticostriatal excitatory circuit, predominantly supported by the PT→ChIs→glutAx→SPNs motif, running in parallel to the canonical excitatory monosynaptic connection from the cortex to SPNs. The photoactivation of that motif evokes a second phase of excitation onto SPNs, mediated by acetylcholine-induced activation of nicotinic receptors at presynaptic glutamatergic terminals in the DLS, and resulting in biphasic corticostriatal signals (**Figure 5K**). Such signals are preferentially evoked by PT, rather than by IT activation, due to a stronger PT→ChIs relative input strength that more efficiently recruits ChIs (**Figure 5K**). In summary, this study dissects, for the first time, the IT and PT corticostriatal connectivity to ChIs, unraveling a circuit motif that robustly elicits ChIs activation upon PT putative motor command signals. Therefore, the results presented here provide new insights on the polysynaptic impact of corticostriatal IT and PT signals onto SPNs, with potential implications for movement, plasticity and learning.

The corticostriatal connection has been extensively studied (**Yeterian and Van Hoesen**, 1978; **Flaherty and Graybiel**, 1993; **Wall et al.**, 2013; **Guo et al.**, 2015; **Hintiryan et al.**, 2016; **Hooks et al.**, 2018; **Johansson and Silberberg**, 2020). However, this connection was usually assumed homogeneous, neglecting the differences between the diversity of cortical inputs impinging onto striatal SPNs and interneurons. In that sense, previous functional studies using electrical or optogenetic stimulation of corticostriatal fibers and testing its impact onto SPNs, would have predominantly recruited the denser bilateral IT pathway, occluding the details of the sparser PT→SPN connection described here. We overcame that limitation by using mouse lines allowing population-specific control of axonal spiking (**Gerfen et al.**, 2013). We found that, in parallel to the direct IT/PT→SPN connection (**Kress et al.**, 2013), both IT and PT pathways contact ChIs with different relative strength, indicating target selectivity of corticostriatal pathways. Interestingly, a recent study testing ipsi- and contralateral striatal innervation from motor and somatosensory cortices showed that ChIs are exclusively contacted by ipsilateral axons, most likely reflecting predominant PT inputs. In that same study, FSIs, the canonical striatal feed-forward inhibitory interneurons (**Mallet et al.**, 2005; **Gittis et al.**, 2010) were proven highly innervated by bilateral corticostriatal fibers, indicating strong IT connectivity (**Johansson and Silberberg**, 2020). Such observation is suggestive of, at least, some degree of corticostriatal IT→FSIs and PT→ChIs specificity, which is partially demonstrated here by our result showing stronger PT relative input strength to ChIs (**Figure 3I**). Although further studies are necessary, this idea becomes especially relevant in the context of a model where ITs are preparatory and PTs broadcast the action command to many motor-related structures (**Li et al.**, 2015). It is, then, tempting to speculate that ITs may permit action preparation by triggering up-states onto action-specific SPNs while silencing action-unrelated SPNs through FSIs. Subsequent PT signals might be key for transitioning from up-states to spikes in the action-related SPNs commanding execution. In this scenario, the feed-forward excitation exerted by PT→ChIs→glutAx→SPNs, by boosting postsynaptic membrane depolarization and input integration, could help securing SPNs spiking and downstream information flow. A possible reason why this second excitatory phase was not detected in the scarce functional studies that have tested PT→SPN connectivity to date, may be on cortical neurons heterogeneity. Recent studies showed that both IT and PT populations could be genetically subdivided into several sub-classes with different synaptic targets (**Economo et al**., 2018; **Tasic et al**., 2018). Thus, since in those cases focal subcortical injections of retrogradely labeling virus were used to achieve PT expression of ChR2 (**Kress et al.**, 2013), it is likely that in previous studies, only a subset of the neurons labelled in the PT-ChR2-EYFP mice were recruited by photostimulation. A similar scenario may underly the variable shape of the EPSCs we evoked when driving ChR2 expression with a viral injection in the cortex (**Figure 1P**). In any case, more research will be required to dissect which PT sub-classes are indeed preferentially targeting ChIs. Understanding the intrastriatal connectivity of PT sub-classes would, for instance, reveal whether during the second EPSC phase, acetylcholine enhances glutamate release from the same axons driving ChIs, or it acts onto other long-range afferents reaching DLS. In fact, exploring the selectivity of acetylcholine for gating different populations of presynaptic long-range axons innervating the DLS, will help elucidating whether PT, IT, thalamic or other presynaptic inputs, alone or in specific combinations, underlie the second phase of excitation reported here. Distinguishing between these scenarios would have strong implications for addressing corticostriatal computations.

Besides driving feed-forward excitation, the activation of the PT→ChIs→glutAx→SPNs motif could gate a window for inducing long-lasting synaptic changes specifically at the activated Cx→SPN contacts, by increasing local levels of acetylcholine. Changes in acetylcholine concentrations are believed to determine the occurrence and the sign (potentiation or depression) of long-term plasticity upon some pre- and postsynaptic activation combinations through muscarinic receptors (**Centonze et al.**, 1999; **Calabresi et al.**, 2000; **Shen et al.**, 2005) and by modulating dopamine levels (**Wang et al.**, 2006; **Shen et al.**, 2008; **Threlfell et al.**, 2012). Dopamine has long been implicated as a key determinant of corticostriatal plasticity onto SPNs (**Shen et al.**, 2008; **Yin et al.**, 2009; **Shan et al.**, 2014; **Yagishita et al.**, 2014) and acetylcholine plays a crucial role in the striatum by modulating the local release of dopamine through nicotinic receptors, independently of distant somatic spiking (**Threlfell et al.**, 2012). Thus, ChIs sit in a strategic position to orchestrate the events underlying corticostriatal plasticity onto SPNs (**Calabresi et al.**, 2000; **Deffains and Bergman**, 2015). In this manner, the PT→ChIs→glutAx→SPNs motif described here may provide a circuit mechanism supporting the delivery of a permissive signal for corticostriatal plasticity onto action-specific SPNs (**Deffains and Bergman**, 2015), conveyed by temporo-spatially restricted changes in striatal levels of acetylcholine, glutamate and maybe dopamine when movement is executed. In fact, the activation of ChIs silence neighboring ChIs by feed-back inhibition, further coordinating spatiotemporal acetylcholine fluctuations (**Sullivan et al.**, 2008; **Dorst et al.**, 2020). Therefore, our work supports a model where PT long-range axons, besides directly selecting or invigorating the execution of a specific action by recruiting SPNs encoding for that action, could also gate plastic changes in corticostriatal synapses onto those SPNs by selectively activating specific ChIs.

An open question from our data is why PT→SPNs fails to evoke spiking in normal ACSF (**Figure 5**)? This happens because the depolarizations evoked by the activation PT inputs alone do not reach spiking threshold in SPNs. Dissimilar expression levels of ChR2 between the transgenic lines could account for this divergence. This technical limitation could coexist with, and it is hard to distinguish from, differences in the IT/PT connectivity strength due to biological constrains that are technically challenging to measure, like, for example, the absolute number of afferents that an individual SPNs sample from each pathway. Another explanation could be the location and distribution of synaptic inputs onto the dendrites of SPNs. It is well established that SPNs need to transition from down-to up-states before spiking (**O’Donnell and Grace**, 1995; **Wilson and Kawaguchi**, 1996). It is also clear that up-states require a precise spatio-temporal coordination of synaptic inputs to occur, with a minimum number of clustered inputs coactivated in a distal dendritic fragment (**Plotkin et al.**, 2011). Thus, one possibility is that IT, but not PT, inputs are arranged onto SPNs in a configuration that could trigger state transitions by themselves. Maybe, in low-magnesium and high-calcium ACSF, boosted PT→ChI activation triggers acetylcholine-dependent glutamate release from different long-range axons innervating SPNs, including IT inputs (**Figure 5K**), with a spatial arrangement that facilitates spiking. Future experiments mapping the organization of corticostriatal inputs onto SPNs will help further elucidating this issue.

In conclusion, our work dissects, for the first time, the IT and PT corticostriatal connectivity to ChIs, uncovering a circuit motif that elicits ChIs activation upon PT putative motor command signals. Altogether, our results propose a model where PT long-range axons, besides directly selecting or invigorating the execution of a specific action by recruiting SPNs encoding for that action, gate corticostriatal plasticity in specific synapses onto those SPNs. Therefore, the results presented here provide new insights on the polysynaptic impact of corticostriatal IT and PT signals onto SPNs, with potential implications for movement, plasticity and learning. Further studies investigating how other striatal microcircuit players sample and integrate long-range IT and PT inputs, as well as pathway-specific roles in vivo, will help tackle these complex corticostriatal circuits underlying motor learning and behavior.

## Acknowledgments

We thank C. Carvalho and A. Vaz for help with the animal care and genotyping. We thank D. Pereira and A. Klaus for comments on the manuscript and members of the Costa Lab for useful discussion. This work was supported by a fellowship/contract from Fundação para a Ciência e a Tecnologia to N.A.M., a European Research Council Consolidator Grant (COG 617142) to R.M.C., and by the Champalimaud Foundation.

## Author Contributions

N.A.M. designed, performed and analyzed the experiments with input from I.I. and R.M.C. A.F.I. did the immunostainings, confocal imaging and neuronal reconstructions with supervision from N.A.M. N.A.M. wrote the paper with input from I.I. and R.M.C.

## STAR Methods

### RESOURCE AVAILABILITY

#### Lead Contact

Further information and requests for resources and reagents should be directed to and will be fulfilled by the Lead Contact, Rui M. Costa.

#### Materials Availability

This study did not generate new unique reagents.

#### Data and Code Availability

The datasets generated during this study are available from the corresponding author on request.

### EXPERIMENTAL MODEL AND SUBJECT DETAILS

#### Animals

All procedures followed the Champalimaud Center for the Unknown Ethics committee guidelines, approved by the Portuguese Veterinary General Board (Ref. No. 0421/000/000/2014). Both male and female transgenic mice ranging from 40 to 76 days of age were used. Mice were allocated to their experimental groups according to their genotype and age, so the age of the recorded neurons from IT and PT cohorts at each experimental condition resulted balanced (**Supplemental Table 1**). Animals were group-housed on a 12 hr light/dark cycle with ad libitum access to food and water. IT-cre (Tlx3 line, STOCK Tg(Tlx3-cre)PL58Gsat/Mmucd, RRID:MMRRC_036670-UCD, GENSAT, http://www.gensat.org/), PT-cre (OE25 line, STOCK Tg(Chrna2-cre)OE25Gsat/Mmucd, RRID:MMRRC_036502-UCD, GENSAT) and ChAT-Cre (B6;129S6^Chattm2(cre)Lowl^/J, Jackson Laboratory, #006410) were crossed with a cre-dependent ChR2-EYFP line (Ai32, B6;129S-Gt(ROSA)26Sor^tm32(CAG-COP4*H134R/EYFP)Hze^/J, Jackson Laboratory, #012569) generating double transgenic lines expressing ChR2-EYFP in specific neuronal populations. In some cases, triple transgenic mice were used by crossing IT- or PT-cre lines with Ai32 line and a transgenic BAC Drd1a-tdTomato line (B6.Cg-Tg(Drd1a-tdTomato)6Calak/J, Jackson Laboratory, #016204). All lines were on C57BL/6 background by backcrossing with C57BL/6J inbred mice for at least 8 generations.

The mouse strains used for this research project, STOCK Tg(Tlx3-cre)PL58Gsat/Mmucd, RRID:MMRRC_036670-UCD and STOCK Tg(Chrna2-cre)OE25Gsat/Mmucd, RRID:MMRRC_036502-UCD’ were obtained from the Mutant Mouse Resource and Research Center (MMRRC) at University of California at Davis, an NIH-funded strain repository, and was donated to the MMRRC by Nathaniel Heintz, Ph.D., The Rockefeller University, GENSAT and Charles Gerfen, Ph.D., National Institutes of Health, National Institute of Mental Health.

### METHOD DETAILS

#### Slice preparation

Mice from 40 to 76 days of age were decapitated after deeply anesthetized with isoflurane. Brains were then dissected in ice-chilled choline chloride solution (110 mM choline chloride, 25 mM NaHCO3, 25 mM d-glucose, 11.6 mM sodium ascorbate, 7 mM MgCl2, 3.1 mM sodium pyruvate, 2.5 mM KCl, 1.25 mM NaH2PO4 and 0.5 mM CaCl2, bubbled with 95% O2/5% CO2) and coronal slices were cut (300 μm thickness) using a Leica VT1200S vibratome. Slices were incubated at 37°C in ACSF (127 mM NaCl, 25 mM NaHCO3, 25 mM d-glucose, 2.5 mM KCl, 2 mM CaCl2,1 mM MgCl2 and 1.25 mM NaH2PO4, bubbled with 95% O2/5% CO2) for 30 minutes before starting recording.

#### Electrophysiology and photostimulation

Data was recorded using a Multiclamp 700B amplifier (Molecular Devices), digitized with a Digidata 1440 (Molecular Devices), and acquired at 10 kHz with pClamp 10 software (Molecular Devices). Neurons were recorded using borosilicate pipettes (resistance 3-5 MΩ, Harvard apparatus) filled with internal solution containing: 135 mM potassium gluconate, 10 mM sodium phosphocreatine, 10 mM HEPES, 3 mM sodium l-ascorbate, 4 mM MgCl2, 4 mM Na2ATP, 0.4 mM Na2GTP and 0.025 mM Alexa-594 (Molecular Probes); pH 7.2; 290 mOsm. In ChI-SPN experiments, biocytin (0.2%, Sigma-Aldrich) was added to the internal solution for neuronal reconstructions. For current-clamp experiments in low-magnesium/high-calcium ACSF, the internal solution was supplemented with Fluo-4 (200 μM, Invitrogen) for other imaging purposes. All recordings were performed in heated (37°C) normal ACSF or low-magnesium/high-calcium ACSF (127 mM NaCl, 25 mM NaHCO3, 25 mM d-glucose, 2.5 mM KCl, 4 mM CaCl2 and 1.25 mM NaH2PO4) perfused at 1.5 – 2 mL/min rate. In voltage-clamp experiments, SPNs and ChIs were clamped at −80 mV and −55 mV, respectively. All recorded neurons were at least at 40 μm depth from the slice surface. For each photostimulation trial, input resistance was monitored with a hyperpolarizing test pulse. After each experiment, SPNs identity was confirmed by imaging their morphology and dendritic spines (filled with Alexa 594, **Supplemental Figure 1A**) at 60x with a BX61WI Olympus microscope, with galvanometer-based scanning system (Bruker) and a 2-photon Ti:sapphire laser (820 nm for imaging Alexa 594, Coherent), controlled by PrairieView software (Bruker). Somatic area and distance between ChIs and SPNs within the same neuronal pair were measured from these images using ImageJ/Fiji (NIH). ChIs resting membrane potential was measured at break-in. Metrics from ChIs spikes (inter-spike interval and half-with) and sag were extracted from current-clamp recordings by running a set of 15 square current steps (1 s duration, 20 pA increase, starting at −160 pA) before bath application of 4-AP and TTX. The position of the recorded neurons in the DLS was visually confirmed at 10x. All neurons were recorded from left DLS.

Wide-field photostimulation was performed using two fiber-coupled ~460 nm LEDs (Doric lenses) attached to the 60x microscope objective (~90° apart from each other), to standardize the distance from the light source to the slice across experiments. In order to maximize the amount of light reaching the slice below the objective, the intensity of the illumination was fixed at maximum power (~10 and ~14 mW for each LED) with a SLC-SA/SV/AA/AV series LED controller (Mightex, Canada). Using PrairieView software, the light pulse duration was varied to elicit responses spanning the range of EPSC/P amplitudes explored. For laser-photostimulation of inputs to SPN-ChI pairs, light was delivered using a Point-Photoactivation module (Bruker) coupled to a 1-photon 473 nm laser (Coherent). The grid was positioned so the soma of the recorded neuron located in the center of a horizontal line placed at one-thirds of the total heigh (**Figure 3D**). Each grid location (spacing, 55 μm) was typically photostimulated twice, following a non-neighboring pattern, maximizing the distance between consecutive laser pulses and resulting in an interval of >6 min between illuminations in the same location (inter-location interval: 30 s; inter-grid repetition interval: 1 min). At the beginning of each grid experiment, laser intensity and duration were calibrated with PrairieView software so that the total input charge for IT→ChIs and PT→ChIs EPSCs were comparable (**Results and Figure 3E and G**).

The current-voltage curve of EPSC 2^nd^ peak experiment (**Supplemental Figure 1C and D**), was recorded in normal ACSF with PTX and using cesium-based internal solution (132 mM Cs-gluconate, 10 mM sodium phosphocreatine, 10 mM HEPES, 3 mM sodium l-ascorbate, 4 mM MgCl2, 4 mM Na2ATP, 0.4 mM Na2GTP).

#### Data analysis

Electrophysiology traces were analyzed using Matlab (MathWorks). Total EPSC/P charge/area was calculated as the integral of the trace in a window of 65 ms starting with photostimulation. EPSC/P 1^st^ peak amplitude was computed as the minimum (EPSC) / maximum (EPSP) value in a window between the start of illumination and 35 ms. To minimize confounding EPSC/P 2^nd^ peak with 1^st^ peak in cases where the 2^nd^ peak had higher absolute amplitude than the 1^st^ peak, the interval from the photostimulation until the detected peak was scrutinized. When an earlier a peak (with amplitude > 25% of the detected peak) followed by a valley (with depth > 5% of the detected peak) was present, the earlier value was counted as the EPSC/P 1^st^.

The decay of the EPSC/P 1^st^ phase was subtracted from the EPSC 2^nd^ phase by subtracting a two-term exponential model from the recorded trace. In order to exclude the datapoints corresponding to the detection window of the EPSC/P 2^nd^ phase from the model fit, we used data from two discontinuous periods (50 ms in total). We defined an initial 10 ms period, starting where the EPSC/P 1^st^ peak decayed 10% of its amplitude and finishing with the start of the detection window, and a second 40 ms period beginning immediately after that window. Thus, the 2^nd^ phase detection period was restricted to a fixed-size window spanning from 10 to 32 ms after the EPSC/P 1^st^ peak decayed to 90% of its amplitude. Within this window, EPSC/P 2^nd^ phase peak amplitude and charge/area were computed as the minimum (EPSC) / maximum (EPSP) value and the integral of the subtracted trace, respectively. EPSC/P 2^nd^ phase was detected when the subtracted trace crossed a negative (EPSC) / positive (EPSP) threshold (3-fold the standard deviation of baseline period) within the detection window. EPSC 1^st^ phase charge was calculated by subtracting the EPSC 2^nd^ phase charge from the total EPSC charge. Charge ratio was calculated by dividing the charge of the EPSC 2^nd^ phase over the total EPSC charge. Peak ratio was computed by dividing the EPSC 2^nd^ peak amplitude over the EPSC 1^st^ peak amplitude. In pharmacological experiments, baseline conditions were computed once the amplitude of the EPSC first peak stabilized after an initial ramping up period (**Supplemental Figure 2**). In **Supplemental Figure 2A**, peak amplitude was normalized to the amplitude of the first trial while charge ratio and peak ratio were normalized to the first detected EPSC 2^nd^ phase.

Latencies to EPSC/P 1^st^ and 2^nd^ peak were computed from the start of the illumination pulse. EPSC/P trial-by-trial and neuron-based analyses used the same analysis criteria. Response probability was calculated by dividing the number of detected second phases by the total number of photostimulations in the bin (trial-by trial, **Figure 1F and I**) or SPNs (neuron-based, **Figure 1N and 5D**). Overall spike latency was computed from the start of photostimulation to the peak of the action potential. This period was then sub-divided into early and late sub-periods using the start of the EPSP (detected when the trace crossed a positive threshold of 3-fold the standard deviation of baseline period), as the border between them. Population spike probability and population spike latency were first calculated for individual neurons (number of spikes/total number of trials in the same light condition, and mean latency to spike in the same light condition, respectively) and then averaged across neurons within the same illumination condition. For spike analysis, only traces with baseline membrane potential below (or equal to) −60 mV were included.

Modulation index was calculated using the formula: (X in ACSF – X in Drug) / (X in ACSF + X in Drug). Where X was: **Figure 2**, charge ratio or peak ratio; **Supplemental Figure 3**, EPSC 1^st^ or 2^nd^ phase charge or EPSC 1^st^ or 2^nd^ phase peak amplitude.

For ChAT-ChR2 experiments, EPSC charge was calculated as the integral of the trace in a window of 65 ms starting with photostimulation. EPSC peak amplitude was computed as the minimum value in a window between the start of illumination and 60 ms. Latencies to EPSC peak were counted from the start of the illumination pulse.

For each neuron in 4-AP and TTX experiments in **Figure 3**, EPSCs recorded in the same location were averaged and charge was computed as the integral of the average trace in a window of 45 ms starting with the light pulse. ChI/SPN input ratio was calculated by dividing the sum of all EPSC charges in the ChI over the sum of all EPSC charges in its paired SPN. ChIs sag difference was calculated as the difference between the mean membrane potential from 5 ms at the beginning and 5 ms at the end (800 ms apart) of a 1 s hyperpolarizing current step of −160 pA.

#### Pharmacology

PTX (100 μM, Tocris); MLA (1 μM, Sigma-Aldrich); DNQX (10 μM, Tocris) + APV (50 μM Tocris) and 4-AP (100 μM, Sigma-Aldrich) + TTX (1 μM, Abcam) were bath applied. The effects of the drugs on the responses were tested after 5 minutes of drug recirculation in the recording chamber. From the 10 MLA experiments in **Figure 2**, in 8 cases MLA was added alone; in 2 cases MLA was added together with the nicotinic blocker Mecamylamine (MEC, 100 μM, Sigma-Aldrich). Monosynaptic connectivity was tested after, at least, 10 minutes of 4-AP and TTX recirculation.

#### Confocal imaging

IT- or PT-ChR2-EYFP mice at ~60 days of age were intracardiacally perfused with 4% paraformaldehyde. Brains were removed and coronally sliced (50 μm thickness). Slices were mounted using Mowiol and imaged with a Zeiss LSM 710 confocal microscope.

#### Immunostaining and neuronal reconstruction

Slices with biocytin-filled ChI-SPN pairs were fixed in 4% paraformaldehyde for at least two hours at 4°C. Slices were then rinsed in PBS and incubated with goat anti-ChAT primary antibody (1:2500 to 1:5000, Chemicon/Millipore cat#AB144P) in PBS with 0.4% Triton-X100 and 2% Normal Horse Serum (NHS, Gibco/Thermo Fisher cat#16050-130) for two overnights at 4°C. After several PBS washes, slices were incubated with donkey anti-goat Alexa-405 conjugated secondary antibody (1:1000, Abcam cat# ab175664) and Streptavidin conjugated with Alexa-594 (1:200, Life Technologies cat#16892) in PBS with 0.4% Triton-X100 and 2% NHS for two hours at room temperature. Slices were mounted using Mowiol and imaged using a Zeiss LSM 710 confocal microscope. Colocalization of Streptavidin and ChAT was assessed using stacks of individual confocal planes at 20x. Dendritic morphology was reconstructed and measured from tiles of 25x images using neuTube (**Feng et al.**, 2015) and Simple Neurite Tracer (**Longair et al.**, 2011) plugin from ImageJ/Fiji (NIH). Reconstruction traces in **Figure 3** were exported using the HBP Neuron Morphology Viewer (**Bakker et al.**, 2017).

#### Stereotaxic surgery

For viral injection in **Figure 1O-P**, a female PT-cre mouse of 24 days was anesthetized with isoflurane (1%–3%, plus oxygen at 1-1.5 l/min) and head-fixed using a stereotaxic frame (David Kopf Instruments, Model 962LS) over a heating pad (ATC1000, World Precision Instruments) at 35-37°C. A small craniotomy was drilled over the left M1 following the coordinates (from bregma): 0.55 mm anterior / 1.5 mm lateral. A pulled glass capillary (Drummond Scientific, USA) with a beveled tip of ~20 μm size was then lowered until reaching 750 μm of depth from the brain surface. 300 nL of AAV5-EF1a-DIO-hChR2(H134R)-EYFP-WPRE-pA (University of North Carolina Vector Core, USA) were then delivered with a Nanoject II Injector (Drummond Scientific, USA) at 4.6 nL/5 s rate. The capillary was removed eight minutes after the last injection pulse. The skin was closed using Vetbond tissue adhesive (3M, USA). The mouse was placed over a heating pad and fully recovered from anesthesia before returning to its homecage.

### QUANTIFICATION AND STATISTICAL ANALYSIS

All statistical analyses were done with Matlab (MathWorks). Since normality was not assumed or rejected using Kolmogorov-Smirnov test, non-parametric two-tailed Wilcoxon rank sum test or two-tailed Wilcoxon signed-rank test were used across the manuscript for independent and paired comparisons, respectively. Statistical significance was defined as: ns, Non-significant, * p < 0.05, ** p < 0.01, *** p < 0.001 and **** p < 0.0001. Sample size were not predetermined but our groups are in line with previous studies (**Tanimura et al.**, 2019). Sample size, specific statistical test, exact p value and additional information are detailed in the figure legends or Results. Data is presented as mean ± standard error of the mean (SEM).

### KEY RESOURCES TABLE

| REAGENT or RESOURCE | SOURCE | IDENTIFIER |
| --- | --- | --- |
| **Antibodies** |  |  |
| Goat anti-ChAT primary antibody | Chemicon/Millipore | Cat#AB144P |
| Donkey anti-goat Alexa-405 conjugated secondary antibody | Abcam | Cat#ab175664 |
| Streptavidin conjugated with Alexa-594 | Life Technologies | Cat#16892 |
| **Bacterial and Virus Strains** |  |  |
| AAV5-EF1a-DIO-hChR2(H134R)-EYFP | University of North Carolina Vector Core | Cat#20298 |

| Chemicals, Peptides, and Recombinant Proteins |  |  |
| --- | --- | --- |
| Alexa Fluor™ 594 dye | Molecular probes | Cat#A10438 |
| Biocytin | Sigma-Aldrich | Cat#B4261; CAS: 576-19-2 |
| Picrotoxin (PTX) | Tocris | Cat#1128; CAS: 124-87-8 |
| Methyllycaconitine citrate salt (MLA) | Sigma-Aldrich | Cat#M168; CAS: 112825-05-5 |
| 6,7-Dinitroquinoxaline-2,3-dione disodium salt (DNQX) | Tocris | Cat#2312; CAS: 1312992-24-7 |
| DL-2-Amino-5-phosphonopentanoic acid sodium salt (APV) | Tocris | Cat#3693; CAS: 1303993-72-7 |
| Tetradotoxine citrate (TTX) | Abcam | Cat#ab120055; CAS: 18660-81-6 |
| 4-Aminopiridine (4-AP) | Sigma-Aldrich | Cat#A78403; CAS: 504-24-5 |
| Mecamylamine hydrochloride (MEC) | Sigma-Aldrich | Cat#M9020; CAS: 826-39-1 |
| Experimental Models: Organisms/Strains |  |  |
| Tg(Tlx3-cre)PL58Gsat/Mmucd (Tlx3, IT-cre) | GENSAT | RRID:MMRRC_036670-UCD |
| Tg(Chrna2-cre)OE25Gsat/Mmucd (OE25, PT-cre) | GENSAT | RRID:MMRRC_036502-UCD |
| B6;129S6<sup>Chatm2(cre)Lowl</sup>/J (ChaT-cre) | The Jackson Laboratory | Cat#006410 |
| B6;129S-Gt(ROSA)26Sor<sup>tm32(CAG-COP4*H134R/EYFP)Hze</sup>/J (Ai32, Chr2-EYFP) | The Jackson Laboratory | Cat #012569 |
| B6.Cg-Tg(Drd1a-tdTomato)6Calak/J | The Jackson Laboratory | Cat #016204 |
| Software and Algorithms |  |  |
| pClamp 10 | Molecular Devices | http://www.moleculardevices.com/products/software/pclamp.html ; RRID:SCR_011323 |
| ImageJ/Fiji | NIH | https://imagej.net/ ; RRID:SCR_003070 |
| Matlab 2018a | MathWorks | http://www.mathworks.com/products/matlab/ ; RRID:SCR_001622 |
| neuTube | Feng et al. 2015 | https://www.neutracing.com/ |
| Simple Neurite Tracer | Longair et al. 2011 | http://imagej.net/Simple_Neurite_Tracer ;<br>RRID:SCR_016566 |
| HBP Neuron Morphology Viewer | Bakker et al. 2017 | https://neuroinformatics.nl/HBP/morphology-viewer/ |

