## Supplemental information for "Pyramidal tract neurons drive feed-forward excitation of striatum through cholinergic interneurons"

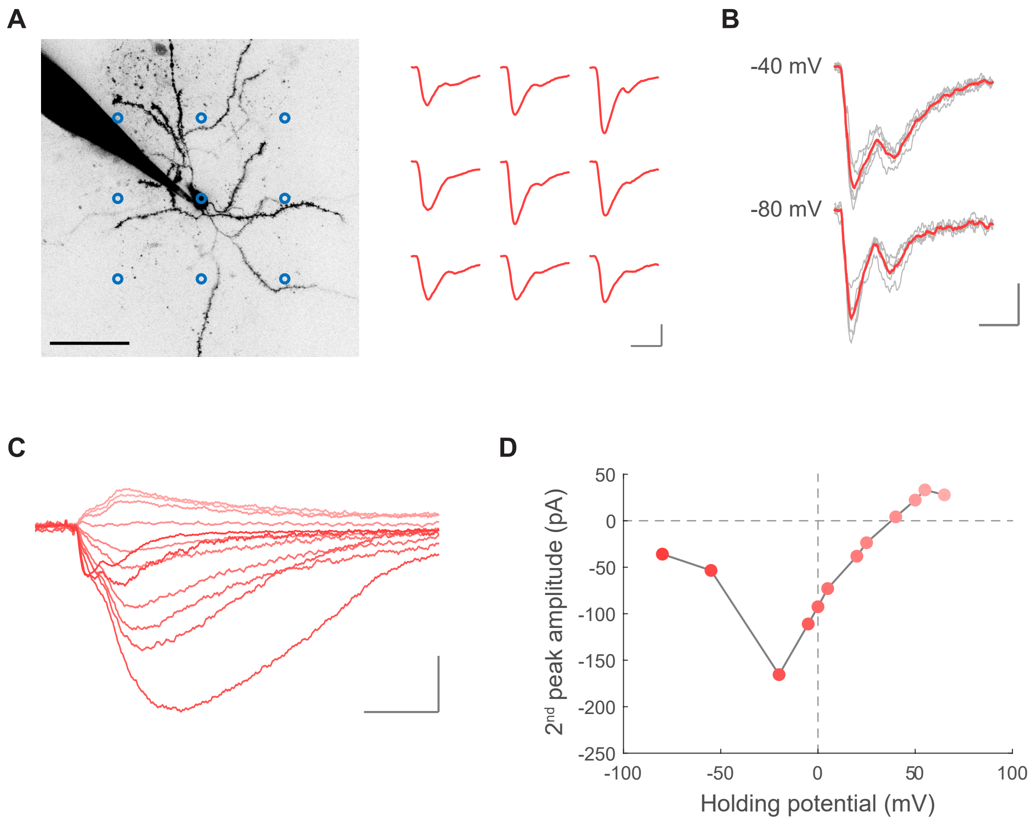


**Supplemental Figure 1. PT→SPN 2nd peak behaves like a cationic current.**

(A) Left, 2-photon image of a SPN showing the grid pattern used for focal photostimulation (blue circles). Scale bar: 50 µm. Right, individual EPSCs elicited when photostimulating PT axons at each grid location with a 473 nm 1-photon laser. Scale bar: 20 ms, 100 pA.

(B) Example of EPSCs elicited by the photostimulation of PT fibers while recording from a SPN in ACSF and held at membrane potentials below (-80 mV) and above (-40 mV) the calculated reversal potential for chloride (-75.6 mV). Thin light gray traces are the five individual trials corresponding to the thicker mean traces. Scale bar: 25 ms, 50 pA.

(C) Example traces of EPSCs elicited by the photostimulation of PT fibers while recording from a SPN in PTX and held at increasing membrane potentials (from dark red to light red: -80, -55, -20, -5, 0, 5, 20, 25, 40, 50, 55, 65 mV). Scale bar: 50 ms, 50 pA.

(D) Quantification of the amplitude of the 2nd peaks evoked in C as a function of the different holding potentials. Shade of red from each datapoint matches its respective trace in C.

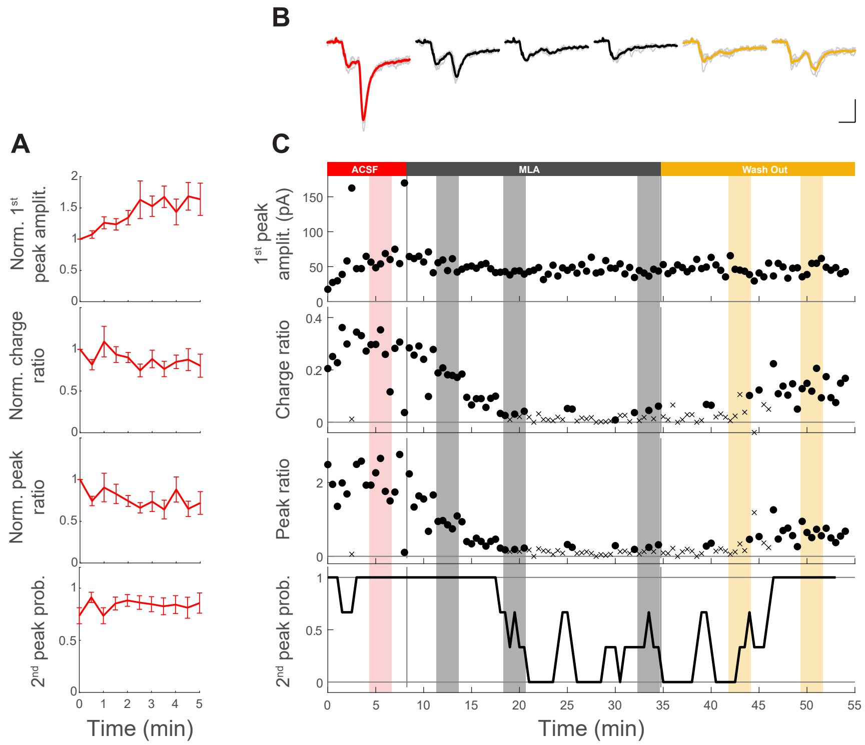


**Supplemental Figure 2. Metrics extracted from PT→SPN EPSCs are reliable within and across experiments.**

(A) From top to bottom: plots showing the normalized amplitude of the 1st phase of the EPSC, the normalized charge ratio, the normalized peak ratio and the response probability for a 2nd peak as a function of time, for all SPNs recorded while activating PT inputs in ACSF. Data is mean ± SEM. For each SPN, data was normalized to its first trial. Each time point is the average of 14-34 SPNs from 19 PT-ChR2-EYFP mice.

(B) Example of an individual experiment showing EPSCs from a SPN when identically photostimulating PT fibers in different bath conditions: ACSF (red), MLA (black), and wash out with ACSF (yellow). Thin light gray traces are the five individual trials underlying the thicker mean traces. Scale bar: 20 ms, 50 pA.

(C) From top to bottom: plots showing the amplitude of the 1st phase of the EPSCs, the charge ratio, the peak ratio and the response probability for a 2nd peak as a function of time, for the whole experiment in B. Data points are individual trials elicited every 30 s (for charge ratio and peak ratio, circles: EPSC with 2nd peak; crosses: EPSC without 2nd peak). Response probability was calculated within a moving window of 3 consecutive trials. Shadowed areas highlight the individual trials (thin light gray traces in B) averaged for each bath condition (red: ACSF; black: MLA; yellow: Wash Out).

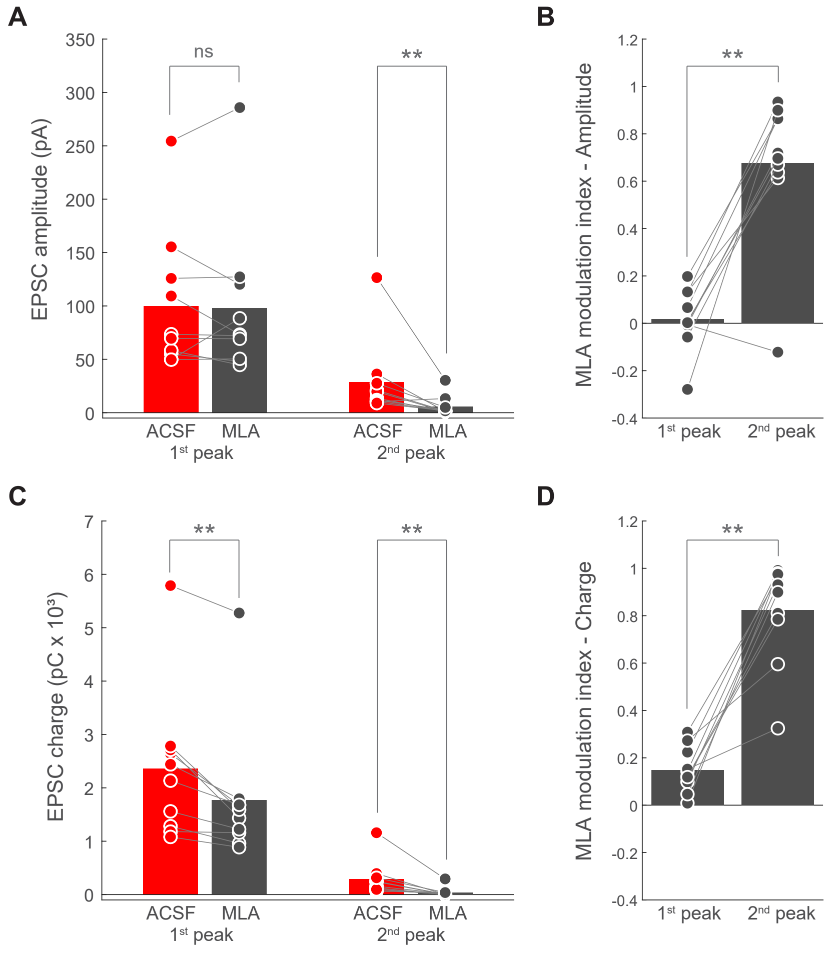


**Supplemental Figure 3. PT→SPN EPSC $1^{st}$ and $2^{nd}$ peak are differentially modulated by nicotinic receptor blocker MLA.**

(A) EPSC amplitude of $1^{st}$ and $2^{nd}$ peak for individual SPNs in ACSF (red) and MLA (black) conditions. n=10 SPNs from 7 PT-ChR2-EYFP mice. Bars represent mean. $1^{st}$ peak p=0.69531; $2^{nd}$ peak p=0.0039063; Wilcoxon signed-rank test.

(B) MLA modulation index of the amplitude of the $1^{st}$ and $2^{nd}$ peak for individual SPNs. n=10 SPNs from 7 PT-ChR2-EYFP mice. Bars represent mean. p=0.0039063; Wilcoxon signed-rank test.

(C) EPSC charge of $1^{st}$ and $2^{nd}$ peak for individual SPNs in ACSF (red) and MLA (black) conditions. n=10 SPNs from 7 PT-ChR2-EYFP mice. Bars represent mean. $1^{st}$ peak p=0.0019531; $2^{nd}$ peak p=0.0019531; Wilcoxon signed-rank test.

(D) MLA modulation index of the charge of the $1^{st}$ and $2^{nd}$ peak for individual SPNs. n=10 SPNs from 7 PT-ChR2-EYFP mice. Bars represent mean. p=0.0019531; Wilcoxon signed-rank test.

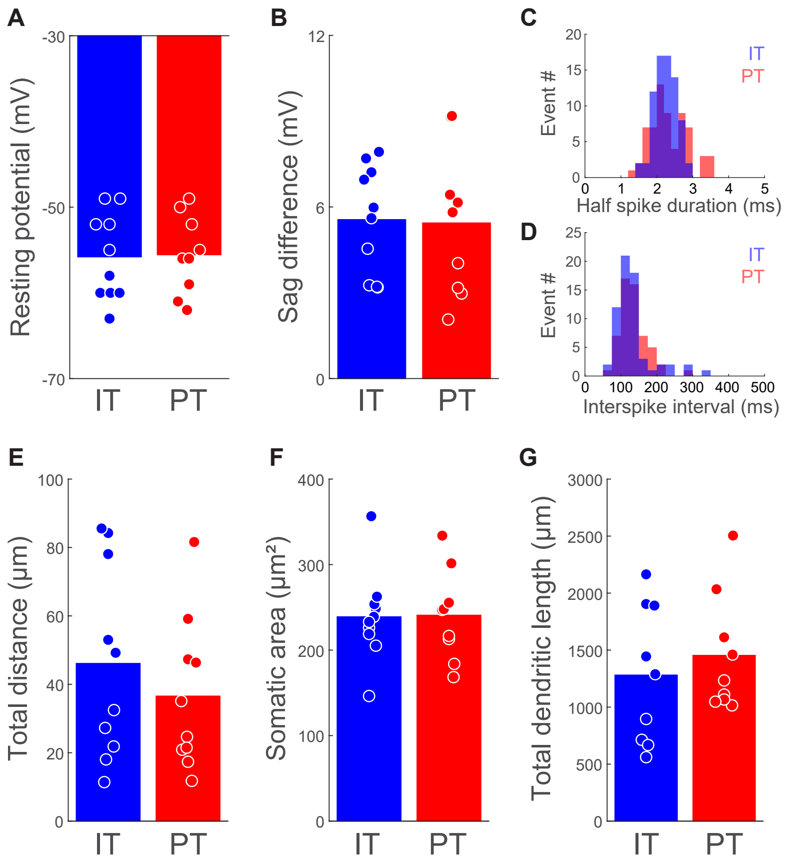


**Supplemental Figure 4. ChIs recorded from IT- and PT-ChR2-EYFP mice belong to a morpho-functional homogenous population.**

(A) Resting membrane potential of ChIs recorded from IT- or PT-ChR2-EYFP mice. Bars represent mean. IT: n=10 ChIs from 10 mice; PT: n=9 ChIs from 9 mice; p=0.9836; Wilcoxon rank sum test.

(B) Difference in membrane potential between the start and the end of the sag evoked upon a 1 s hyperpolarizing current step of -160 pA from ChIs recorded from IT- or PT-ChR2-EYFP mice. Bars represent mean. IT: n=10 ChIs from 10 mice; PT: n=9 ChIs from 9 mice; p=0.7197; Wilcoxon rank sum test.

(C) Histogram showing the distribution of half spike durations upon a 1 s depolarizing pulse of +60 pA from the ChIs recorded from IT- or PT-ChR2-EYFP mice. IT: n=74 spikes from 10 ChIs from 10 mice. PT: n=65 spikes from 9 ChIs from 9 mice; p=0.3982; Wilcoxon rank sum test, z=0.84.

(D) Histogram showing the distribution of inter-spike intervals upon a 1 s depolarizing pulse of +60 pA from the ChIs recorded from IT- or PT-ChR2-EYFP mice. IT: n=64 intervals from 10 ChIs from 10 mice. PT: n=56 intervals from 9 ChIs from 9 mice; p=0.18065; Wilcoxon rank sum test, z=1.34.

(E) Distance between an individual ChI and its paired SPN recorded from IT- or PT-ChR2-EYFP mice. Bars represent mean. IT: n=10 pairs from 10 mice; PT: n=10 pairs from 9 mice; p=0.42736; Wilcoxon rank sum test, z=0.79.

(F) Somatic area of ChIs from IT- or PT-ChR2-EYFP mice. Bars represent mean. IT: n=10 ChIs from 10; PT: n=9 ChIs from 9 mice; p=0.96824, Wilcoxon rank sum test.

(G) Total dendritic length of ChIs recorded from IT- or PT-ChR2-EYFP mice. Bars represent mean. IT: n=9 ChIs from 9 mice; PT: n=9 ChIs from 9 mice; p=0.48943; Wilcoxon rank sum test.

| | | IT age (days) | PT age (days) | Rank sum |
|---|---|---|---|---|
| Figure 1 | Mean ± SEM | 57.00 ± 1.75 | 56.12 ± 1.30 | p = 0.4586 |
| | Range | 43 - 66 | 42 - 65 | z = 0.7412 |
| | n (SPNs) | 26 | 34 | |
| Figure 3 | Mean ± SEM | 65.10 ± 1.55 | 67.40 ± 2.18 | p = 0.2879 |
| | Range | 58 - 72 | 53 - 74 | z = -1.0627 |
| | n (ChI-SPN pairs) | 10 | 10 | |
| Figure 5 | Mean ± SEM | 44.27 ± 0.27 | 44.82 ± 0.82 | p = 0.3812 |
| Normal | Range | 43 - 45 | 41 - 47 | z = -0.8757 |
| ACSF | n (SPNs) | 11 | 11 | |
| Figure 5 | Mean ± SEM | 44.69 ± 0.76 | 44.71 ± 1.55 | p = 0.8091 |
| 0 $Mg^{++}$ / 4 | Range | 40 - 49 | 41 - 53 | z = 0.2416 |
| $Ca^{++}$ ACSF | n (SPNs) | 13 | 7 | |

**Supplemental Table 1. Neuronal age is similar between IT and PT groups within the same experimental condition.**
